# Bioinformatics workflows for genomic analysis of tumors from Patient Derived Xenografts (PDX): challenges and guidelines

**DOI:** 10.1101/414946

**Authors:** Xing Yi Woo, Anuj Srivastava, Joel H. Graber, Vinod Yadav, Vishal Kumar Sarsani, Al Simons, Glen Beane, Stephen Grubb, Guruprasad Ananda, Rangjiao Liu, Grace Stafford, Jeffrey H. Chuang, Susan D. Airhart, R. Krishna Murthy Karuturi, Joshy George, Carol J. Bult

**Affiliations:** The Jackson Laboratory for Genomic Medicine, Farmington, CT 06030, USA; The Jackson Laboratory, Bar Harbor, ME 04609, USA; MDI Biological Laboratory, Bar Harbor, ME 04609, USA

## Abstract

Bioinformatics workflows for analyzing genomic data obtained from xenografted tumor (e.g., human tumors engrafted in a mouse host) must address several challenges, including separating mouse and human sequence reads and accurate identification of somatic mutations and copy number aberrations when paired normal DNA from the patient is not available. We report here data analysis workflows that address these challenges and result in reliable identification of somatic mutations, copy number alterations, and transcriptomic profiles of tumors from patient derived xenograft models. We validated our analytical approaches using simulated data and by assessing concordance of the genomic properties of xenograft tumors with data from primary human tumors in The Cancer Genome Atlas (TCGA). The commands and parameters for the workflows are available at https://github.com/TheJacksonLaboratory/PDX-Analysis-Workflows.

## Introduction

Patient-Derived Xenograft (PDX) models are *in vivo* preclinical models of human cancer for translational cancer research and personalized therapeutic selection [1-7]. Previous studies have demonstrated engrafted human tumors retain key genomic aberrations found in the original patient tumor [3, 8, 9] and that treatment responses of tumor-bearing mice typically reflect the responses observed in patients [6, 10]. PDXs have been used successfully as a platform for pre-clinical drug screens [6, 7, 10], to facilitate the development of potential biomarkers of drug response and resistance [6, 7, 11], and to select appropriate therapeutic regimens for individual patients [8].

The Jackson Laboratory (JAX) PDX Resource has over 400 PDX models from more than 20 different types of cancer. A schematic summarizing the processes used for model generation, quality control, and characterization process for the resource is shown in Figure 1. Genome characterization of PDX tumors includes the identification of somatic mutations, copy number alterations, and transcriptional profiles. Over 100 of the models have been assessed to date for responses to various therapeutic agents. The integration of results from dosing studies with genomic data for the models has been successfully applied to the identification of novel genomic biomarkers associated with treatment responses [12].

**Figure 1.**
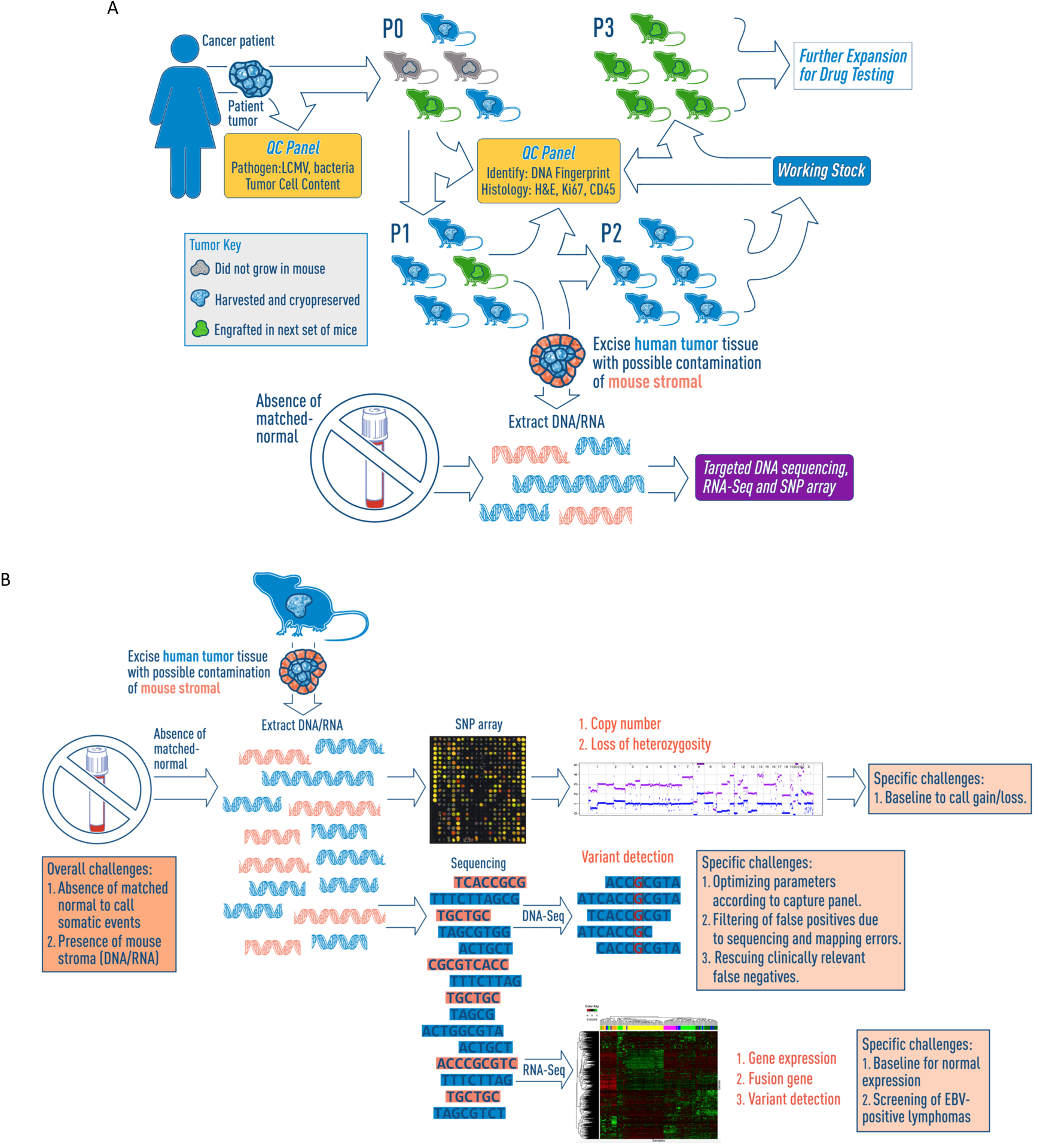
(A) The Jackson Laboratory (JAX) has generated, clinically annotated, and genomically characterized more than 450 patient-derived xenograft (PDX) cancer models from about 20 different types of cancer using the immunodeficient NOD.Cg-*Prkdc*^*scid*^ *Il2rg*^*tm1Wjl*^/SzJ (aka, NSG™) mouse as the host strain (http://tumor.informatics.jax.org/mtbwi/index.do). This figure shows the workflow of PDX model generation from patient tumor, the process of engraftment and passaging that supplies to the JAX PDX resource, and the generation of genomic and transcriptomic data to profile the PDX models. (B) The PDX models are profiled by: 1) DNA mutations from capture sequencing using the JAX Cancer Treatment Profile™ (CTP, https://www.jax.org/clinical-genomics/clinical-offerings/jax-cancer-treatment-profile), the Illumina Truseq panel or whole-exome sequencing, 2) DNA copy-number variations using Affymetrix SNP 6.0 arrays, and 3) gene expression profiles from Affymetrix microarrays or RNA sequencing (Illumina HiSeq). The analysis of the genomic and transcriptomic data of PDX models poses several challenges which we have developed several strategies to circumvent these issues.

To generate accurate calls for mutations and copy number variants for human tumors engrafted in a mouse host, several challenges had to be addressed. First, because human stroma is replaced by mouse cells and tissues during tumor engraftment, sequence data generated for PDX tumors includes both mouse and human sequences. As the protein-coding regions of the mouse and human genomes are 85% identical on average, there is a high risk of introducing false positive variants in functional regions and erroneous gene expression [13-15]. Second, because the tumor material used to create models in the JAX PDX Resource consisted of material that remained following clinical pathology assessment (i.e. material was not collected specifically for xenograft model creation), paired normal samples were not available for the majority of tumor samples used to generate the PDXs. The absence of normal tissue complicates the ability to distinguish germline variants from somatic alterations (point mutations, indels and copy number aberrations) in the tumor [16-19]. Third, false positive (FP) variants due to errors in sequencing and mapping require additional filtering steps in the computational workflow [20-22]. Finally, it has been reported previously that the immunodeficient host mice are susceptible to forming B-cell human lymphomas during engraftment due to Epstein-Barr virus (EBV)-associated lymphomagenesis [23-27]. Systematic screening of PDX tumor samples for EBV transformation is an important step in quality assurance for the integrity of PDX repositories.

Here, we describe bioinformatics analysis workflows and guidelines (https://github.com/TheJacksonLaboratory/PDX-Analysis-Workflows) that we developed for the for the analysis of genomic data generated from PDX tumors (http://www.tumor.informatics.jax.org/mtbwi/pdxSearch.do). These workflows incorporated established tools and public databases and were tailored to address the specific challenges mentioned above by tuning parameters and addition of filters. We demonstrate how our methods, using simulated and experimental data, improve the accuracy in the detection of somatic alterations in PDX models. We also developed a classifier based on expression data to systematically identify and filter out EBV transformed samples. Finally, to verify the effectiveness of our workflows, we show the overall concordance of the genomic and transcriptomic profiles of the PDX models in the JAX PDX resource with relevant tumor types from The Cancer Genome Atlas (TCGA).

## Results

### Workflow for calling somatic point mutations and indels in PDX tumors

A schematic of the variant calling workflow we implemented for human tumors engrafted in mice is shown in Figure 2A and 2B (see Methods).

**Figure 2.**
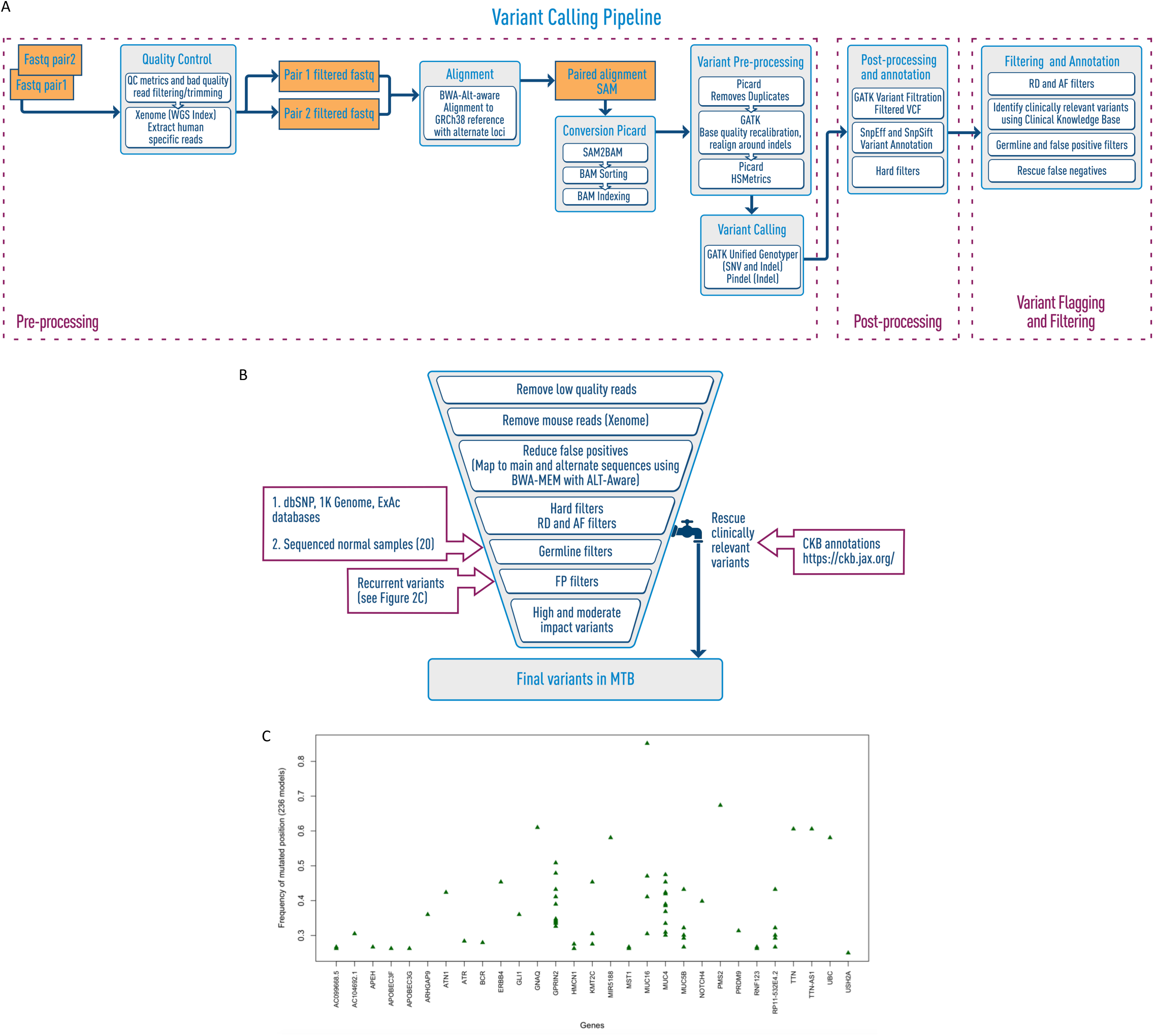
(A) This flow chart describes the variant calling pipeline for PDX DNA sequencing data. (B) This figure shows the different filters used the variant calling pipeline for PDX DNA sequencing data applied to the CTP panel sequencing (see Methods for details). MTB is the Mouse Tumor Biology Database in JAX, PDX models in the JAX PDX resource can be searched in http://tumor.informatics.jax.org/mtbwi/pdxSearch.do. (RD: Read depth, AF: Allele-frequency, FP: False positives (C) The recurrent frequencies of the mutated positions (after germline filtering) for various genes that were found to be recurrent in more than 25% of PDX samples. These were identified as additional false positive variants due to sequencing errors or mapping issues.

#### Preprocessing and removal of mouse reads

Human and mouse DNA reads were classified by Xenome [13], which had shown reliable performance in separate studies [28], and only human reads were used for subsequent variant calling. The percentage of mouse reads within the PDX samples in the JAX resource has a median value of 5.30% (range: 0.00163% - 65.1%) (Figure 3A). Using simulated CTP datasets, we verified that omitting the Xenome step to filter the mouse reads resulted in very low precision (Figure 3B), i.e. large number of FPs, in the absence of the quality hard filters (Supplementary Table S1). These FPs were due to mouse reads being aligned to the reference genome with mismatches and subsequently called as variants with low quality scores (QD).

**Figure 3.**
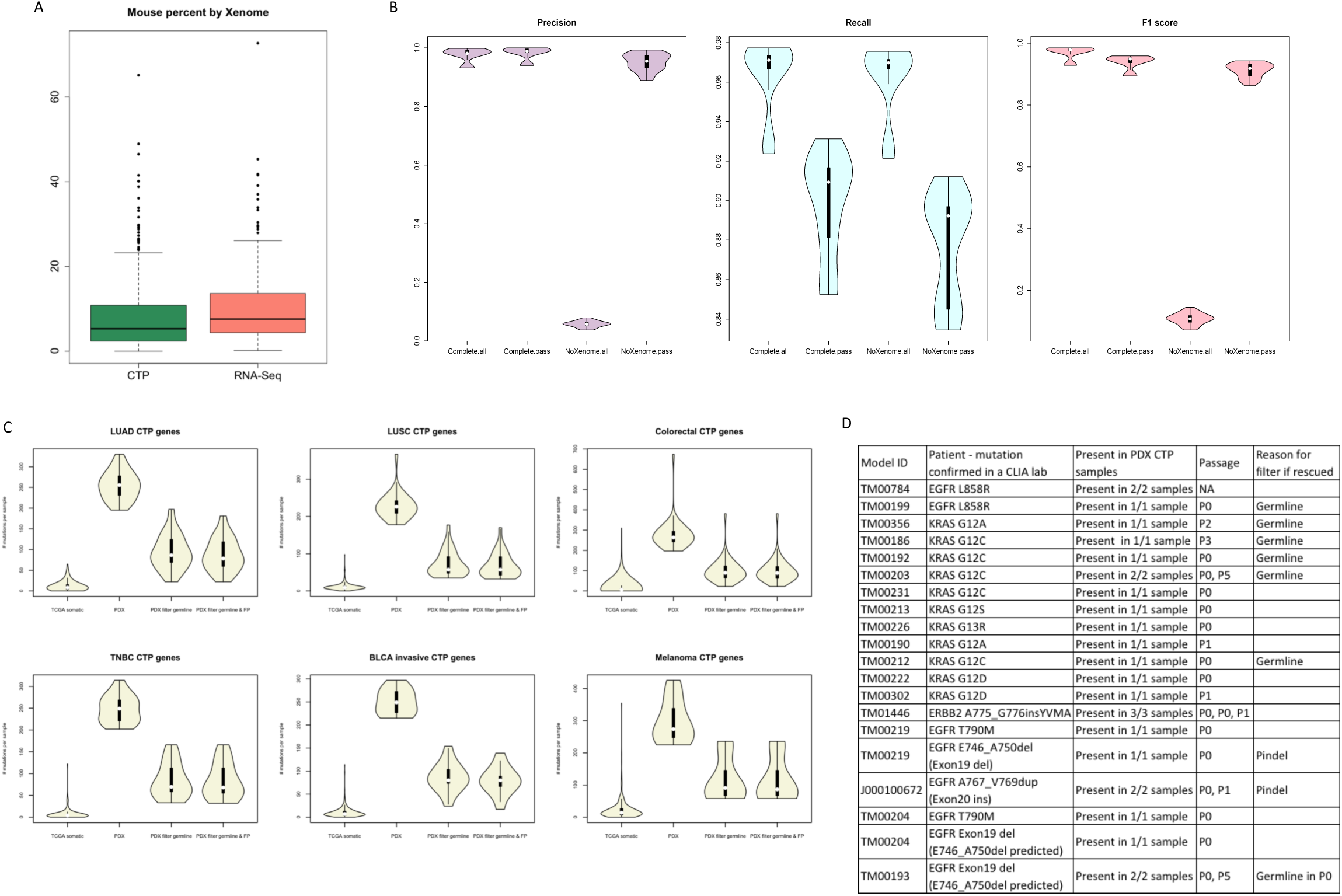
(A) Proportion of mouse reads detected by Xenome for CTP and RNA sequencing data of PDX models. (B) This figure shows the benchmarking of the CTP variant calling pipeline using 45 simulated sequencing datasets different samples, sequencing coverages, and mouse DNA content (see Supplementary Table S2) using precision, recall and F1 score based on the input variants for each sample. Complete: variant calling pipeline with all steps included; NoXenome: variant calling pipeline with Xenome omitted; all: all variants called by the pipeline; pass: variants annotated as “PASS” in the pipeline which pass the hard filters, minimum read depth and minimum alternate allele frequency of the variant. (C) Distribution of mutational load per sample of non-silent coding somatic mutations of CTP genes from exome sequencing TCGA samples and from CTP-panel sequencing of PDX models. TCGA somatic: TCGA somatic mutations reported in maf files; PDX: all variants annotated as “PASS” (pass the hard filters, minimum read depth and minimum alternate allele frequency of the variant); PDX filter germline: all variants annotated as “PASS” and filtered from putative germline variants; PDX filter germline & FP: all variants annotated as “PASS” and filtered from putative germline variants and false positives. (D) Mutations in PDX models that were detected by CTP-panel sequencing and experimentally validated in the corresponding patient tumor. Some of these variants were rescued after initial filtering.

While the default thresholds for GATK hard filtering parameters [29] removed a large proportion of the FPs, applying Xenome to filter for human reads yielded superior performance in terms of substantially higher precision, as well as improvement in recall. In addition, Xenome filtering maintained the correlation between the predicted versus actual allele frequencies, which would otherwise decrease with higher mouse contamination (Supplementary Table S2).

#### Filtering germline variants

To enhance filtering out germline variants from somatic mutations, we sequenced and analyzed 20 normal blood samples using the CTP targeted panel. As shown in Supplementary Figure S1A and S1B, 87% of the variants identified in normal blood had allele frequencies of 40% - 60% or >90% across all the samples, indicating the presence of heterozygous or homozygous common variants, respectively. Ninety-one percent of the variants identified in these 20 samples were annotated in the public germline databases. 4% of these variants were not found in public germline databases, but were recurrent in these normal samples or across the PDX tumors in our collection (Supplementary Figure S1C) and so were added to our list of putative germline variants. Only 5% of all of the variants in the 20 samples were private events. Based on these observations, the variants in each PDX tumor with an allele frequency of 40% - 60% or >90%, and present in either public germline database or our list of putative germline variants (Supplementary Table S3) were filtered out as germline variants (Supplementary Table S3). This was a more conservative approach given that these known germline variants in regions of copy number alterations where the ratio of both alleles were not balanced would not be filtered. Figure 3C shows that the germline filters effectively rectified the estimated somatic mutational load in the PDX tumors (Supplementary Table S5) by about four-fold reduction (Supplementary Table S4), which was reasonable as a large proportion of the variants were expected to be germline.

#### Filtering false positives due to systematic errors

Putative somatic variants with no known effects in cancer that recur across large numbers of PDX samples are potentially FPs arising from sequence assembly based error in the reference the genome, sequencing errors or alignment errors in low mappability regions [30]. To detect these, we filtered out the variants at loci that were recurrently mutated in ≥25% of PDX tumors (Figure 2C). The distribution of tumor types for each of these recurrently mutated positions (n=52) was highly similar to the overall distribution of tumor types in the PDX resource (Supplementary Figure S2A) with Pearson correlation coefficient >0.9 (Supplementary Figure 2B). This implies that these mutations were systematic errors and were not selected for any tumor type, and thus, biologically irrelevant. Filtering these highly recurrent loci did not significantly reduce the predicted mutational load per tumor (Figure 3C and Supplementary Table S4).

#### Rescuing variants

The germline filters might filter out actual somatic events in each PDX sample, leading to false negatives. However, retaining all variants represented in cancer variant databases such as COSMIC would lead to excess FPs. For example, 46% of the variants in the normal samples are present in the COSMIC database (Supplementary Figure S1A). To address the balance of false positive and false negative mutation calls, we “rescued” variants that were initially filtered out based on curated annotations available in the JAX-Clinical Knowledgebase (CKB, https://ckb.jax.org/) [31]. The criteria for rescuing variants included those with 1) known or predicted gain or loss of protein function, 2) potential treatment approach for any cancer type and 3) drug sensitivity and resistance effects in clinical or preclinical studies (Supplementary Table S4). We also included an additional indel caller, Pindel [32], in the workflow in order to increase the sensitivity of indel prediction. As Pindel results contained a large number of FPs, we only included those that were present in the JAX-CKB by the same criteria. Overall, 127 unique variants from 52 genes (1.03% of the total and 2.21 % of the filtered unique variants detected by the CTP platform) were rescued from 381 PDX CTP samples. Nine of these mutations have been validated to be present in the PDX model (Figure 3D). Almost all were initially filtered as germline events, as many well-known actionable cancer mutations (e.g. BRAF V600E and KRAS G12C) are present in the dbSNP database and were filtered if they fall within the germline allele frequency. Two other variants that were not called by GATK initially but were detected by Pindel were rescued as they were annotated clinically relevant.

#### Optimized workflow achieves high performance in somatic mutation calling

Figure 3B shows that our full feature workflow on the simulated datasets achieved the highest precision in variant calling, with insignificant compromise on the recall (Supplementary Table S1). We observed that the allele frequencies of the true positive (TP) variants correlates well (Pearson correlation coefficient >0.99) with the input allele frequencies for all samples (Supplementary Figure S3 and Figure S4, and Supplementary Table S2). Although the estimated allele frequencies were lower than the true allele frequencies, this difference was marginal and could be attributed to the reads carrying the variants being classified as non-human reads by Xenome or not mapped to the genome. Moreover, all (20 out of 20) clinically relevant mutations experimentally validated or clinically reported in the corresponding patient tumors were detected in the PDX tumors (Figure 3D).

### Gene expression analysis in PDXs

A schematic overview of the PDX gene expression workflow is provided in Figure 4A (see Methods).

**Figure 4.**
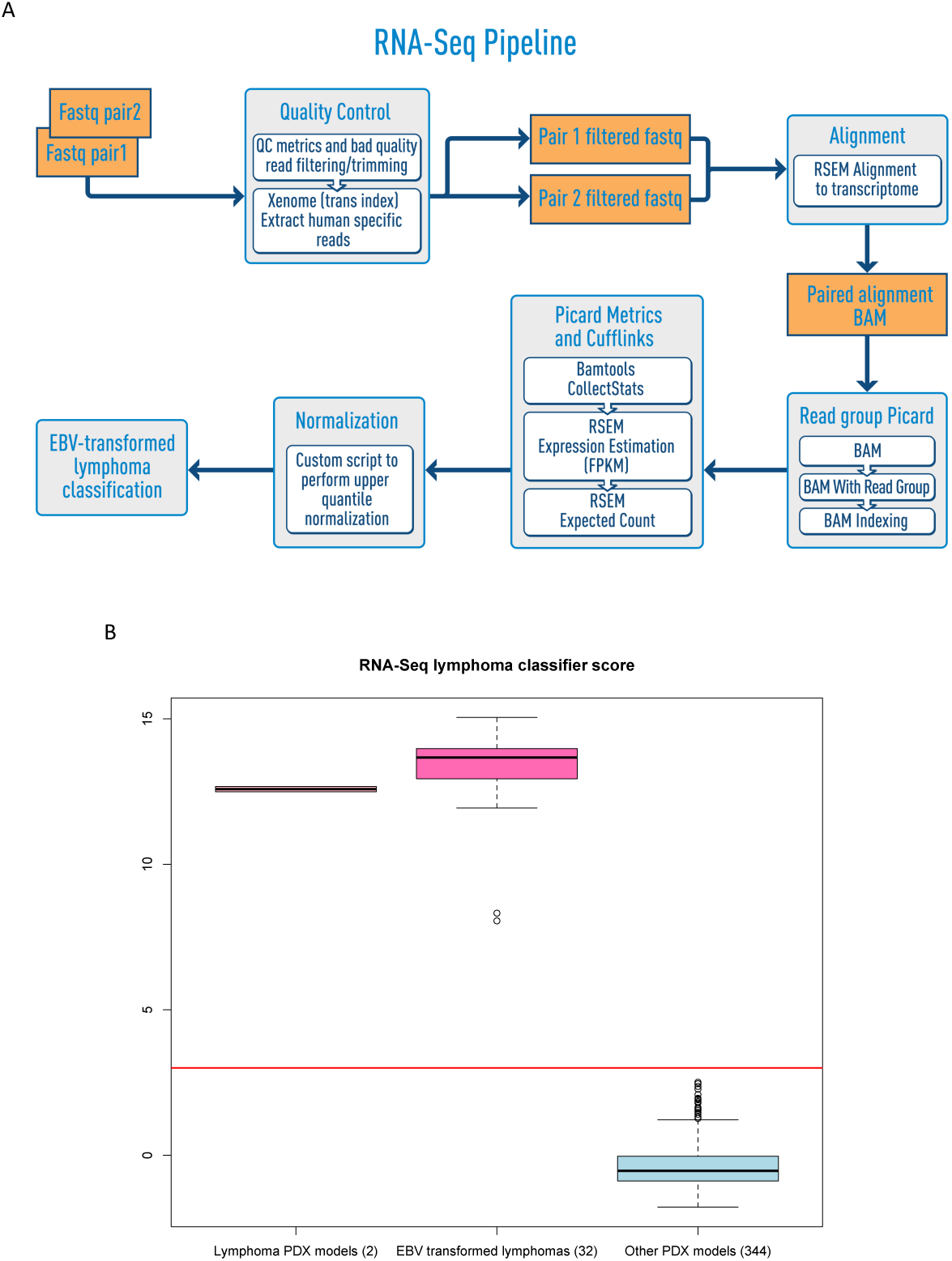
(A) This flow chart describes the RNA expression pipeline and fusion gene prediction for PDX RNA sequencing data. (B) Distribution of lymphoma classification scores of PDX tumors.

#### Screening of EBV-associated lymphomas by RNA-Seq expression data

We observed that the EBV-associated lymphoma tumors that arise in PDX samples display a distinct and highly reproducible expression pattern, regardless of the platforms in which the expression was measured (RNA-Seq, Affymetrix Human Gene 1.0 ST arrays and Human Gene 133 Version 2 arrays). The PDX tumors identified as EBV-associated routinely showed higher correlation in expression profiles than distinct pairs of PDX models derived from common original tumor materials (Supplementary Figure S5). This expression profile was also independent of the tissue of origin of the tumors from which the EBV-associated lymphomas were derived. Given the high similarity in expression profiles, we identified a gene signature based on the most differentially expressed genes between EBV-associated lymphomas and non-EBV-associated tumors (data not shown). Using gene set analysis, we observed that genes associated with B-lymphocytes and other immune processes were over-expressed, while cell-to-cell communication and adherence genes were suppressed (data not shown). We developed a classifier that scored each PDX sample based on the expression levels of the genes in the gene signature (Supplementary Table S6). This single score, when applied on RNA-Seq data, was able to effectively distinguish PDX tumors that were either EBV-transformed or originated from human lymphomas from non-lymphoma PDX tumors (Figure 4B). Overall, 8.5% (32 out of 376) of the non-lymphoma PDX samples with RNA-Seq data in the PDX resource progressed to EBV-associated lymphomas. These tumors were further confirmed to be CD45 positive by immunohistochemistry (IHC) staining, which is the primary tool at JAX to identify PDX tumors that are EBV-transformed.

### Copy Number Variant (CNV) analysis in PDXs

A schematic overview of the PDX CNV workflow is provided in Figure 5A (see Methods).

**Figure 5.**
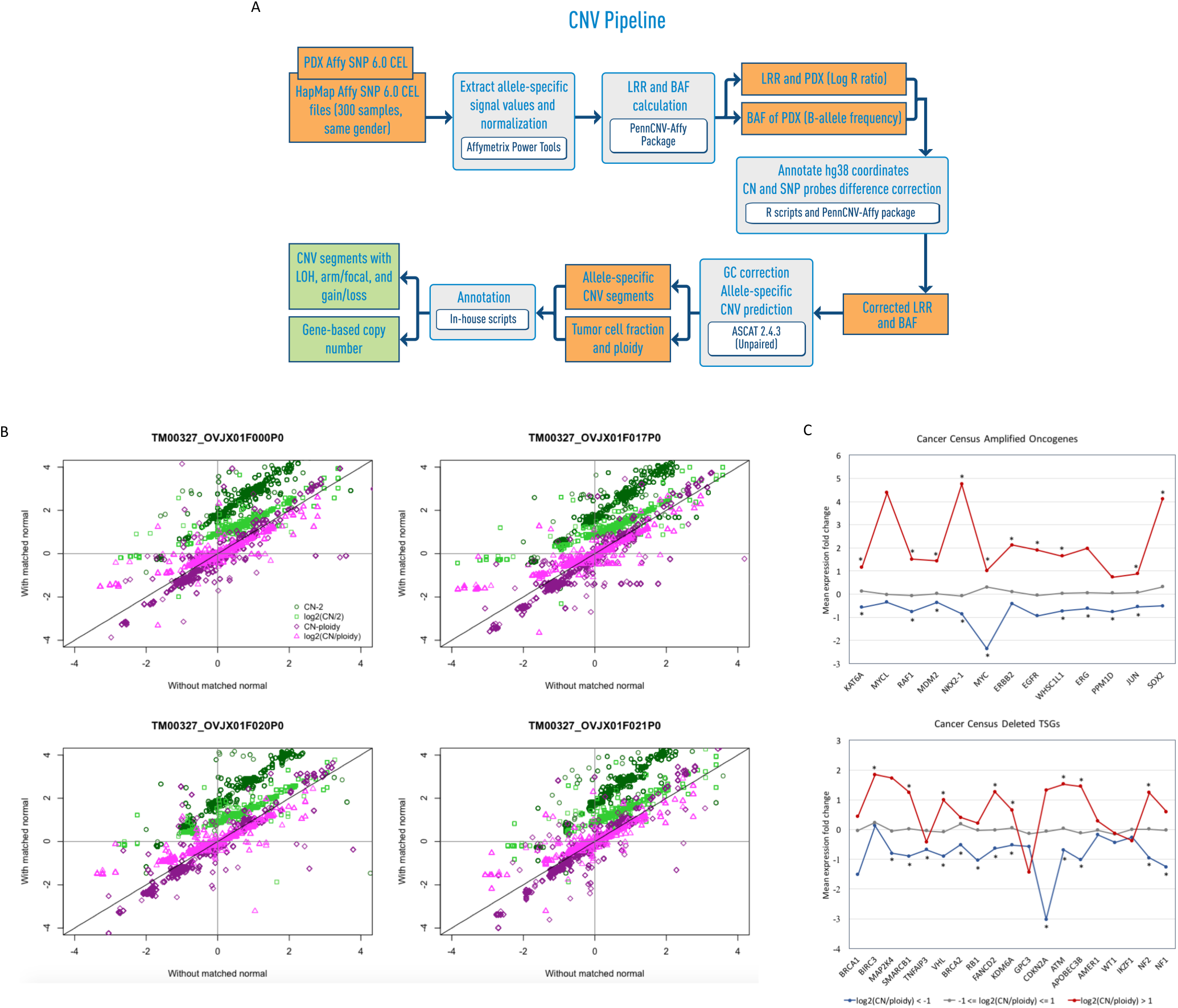
(A) This flow chart describes the CNV and LOH prediction pipeline for PDX SNP array data. (B) Comparison of copy number relative to the estimated overall ploidy of the PDX sample or the diploid state between analyses with and without matched normal. (C) Mean expression fold change of genes with copy number normal, gain and loss state for a selected list of known oncogenes that are amplified in cancers and known tumor suppressor genes that are deleted in cancers from the Cancer Census [34]. Overexpressed and under-expressed genes marked with * indicates significant differences in expression fold change with copy number gain or loss state respectively relative to the normal state across all PDX samples.

#### Effect of mouse DNA on CNV calls

We studied the effect of mouse contamination on array data by hybridizing DNA of the NSG mouse on the human SNP array, and observed that the signal intensity from mouse DNA is negligible (Supplementary Figure S6). Samples with higher mouse content are more likely to result in failure of the standard array quality control or the analysis workflow, due to lower amount of human DNA to give sufficient probe signal, thus enabling samples with substantial mouse contamination to be screened out.

#### Absence of matched normal to call somatic copy number aberrations

We compared the results of the single-tumor CNV analysis with the tumor-normal CNV analysis to access the reliability of the single-tumor CNV analysis results. For the limited number of PDX samples with paired normal samples, we observed overall high similarity between the segmented copy number profiles analyzed with and without the paired-normal sample (Supplementary Figure S7). The gene-based log2(total CN/ploidy) showed good correlation between the single-tumor and tumor-normal CNV analysis (Pearson correlation >0.81, n=9), with 8 out of 9 PDX samples having a correlation of >0.93 (Supplementary Table S7), indicating that the single-tumor CNV analysis was sufficiently robust.

#### Establishing the appropriate baseline to call copy number gains and losses

We analyzed the effects of using different baselines for “normal state” to compute copy number gains and losses using a list of significantly amplified and deleted genes from TCGA (Supplementary Figure S8). When the overall cancer genome ploidy was used as the normal baseline, we observed a balance of a larger proportion of the significantly amplified being called copy number gain, and similarly a larger proportion of the significantly deleted genes being called copy number loss among the PDX samples (Supplementary Figure S9). However, more of both significantly amplified and deleted genes were being classified as amplified when copy number aberrations were calculated relative to the diploid state. While the average ploidy could be estimated differently across the samples for the same model, the copy number changes relative to ploidy remained consistent (Figure 5B and Supplementary Figure S7).

#### Effects copy number aberrations on expression changes

We observed that the estimated copy number gains and losses of known oncogenes (n=23) and tumor suppressor genes (n=40) [33], relative to the average ploidy per PDX sample, generally results in expression fold change (relative to the average expression at copy number normal state) in the same direction (Supplementary Table S8) [11, 34, 35]. Most of these genes show significant over-expression with copy number gain and significant under-expression with copy number loss across the PDX samples (p<0.05) (Figure 5C and Supplementary Figure S10). This shows that the baselines to call copy number gain and loss, and over and under-expression, were correctly established. This significant observation, however, did not hold when we did a global analysis across all genes instead of selected oncogenes and tumor suppressor genes. This was because many genes were not expressed in the respective tissue types even though they were in regions affect by copy number alterations, and the expression of many genes, despite being non-altered regions, could be regulated by other mutations or epigenetic mechanisms in the tumors.

### Comparison of genomic and transcriptomic profiles of PDX models and TCGA patient tumors

Due to the lack of paired-normal samples for the PDX models in the JAX PDX Resource, we were unable to experimentally validate the somatic calls predicted from the various workflows. To determine if the results of our genomic analysis workflows were similar to known somatic profiles of the same tumor type, we compared the overall genomic and transcriptomic profiles for selected tumor types between the JAX PDX resource and patient tumor cohorts in the TCGA.

#### Frequently mutated genes in primary patient tumors in TCGA detected in the PDX resource

The distribution of somatic coding non-silent mutational load of the CTP genes for each tumor type was comparable between PDX and TCGA (Figure 6A). Despite the much smaller sample size for each PDX tumor type, we still observed higher mutational load in colorectal cancer and melanoma. Nonetheless, the overall mutational load remained higher in PDX tumors, which could be possibly due to the fact that the PDX tumors were sequenced at a higher coverage (>900X) using the CTP targeted panel, and thus more variants were detected per base pair compared to exome sequencing (∼100X) of TCGA tumors. Moreover, known germline variants with allele frequency outside the range of 40% - 60% and >90%, possibly due to errors in allele frequency estimation or copy number aberrations at the variant position, as well as private germline variants, were not filtered. The mutations in TCGA were curated with partial experimental validations, hence the mutation count and FP rate were expected to be lower. Given that there were more samples in the TCGA cohorts, we compared the genes that were mutated at 5% frequency with genes that were mutated in at least one sample within the same tumor type in the PDX resource. Almost all genes mutated at high frequencies in TCGA tumors were mutated in PDX tumors, with significant p-values (p < 1×10^−4^) by Fisher’s exact test (Figure 6B, Supplementary Table S9). This indicates that the key drivers by mutation within each cancer type were preserved in PDX tumors.

**Figure 6.**
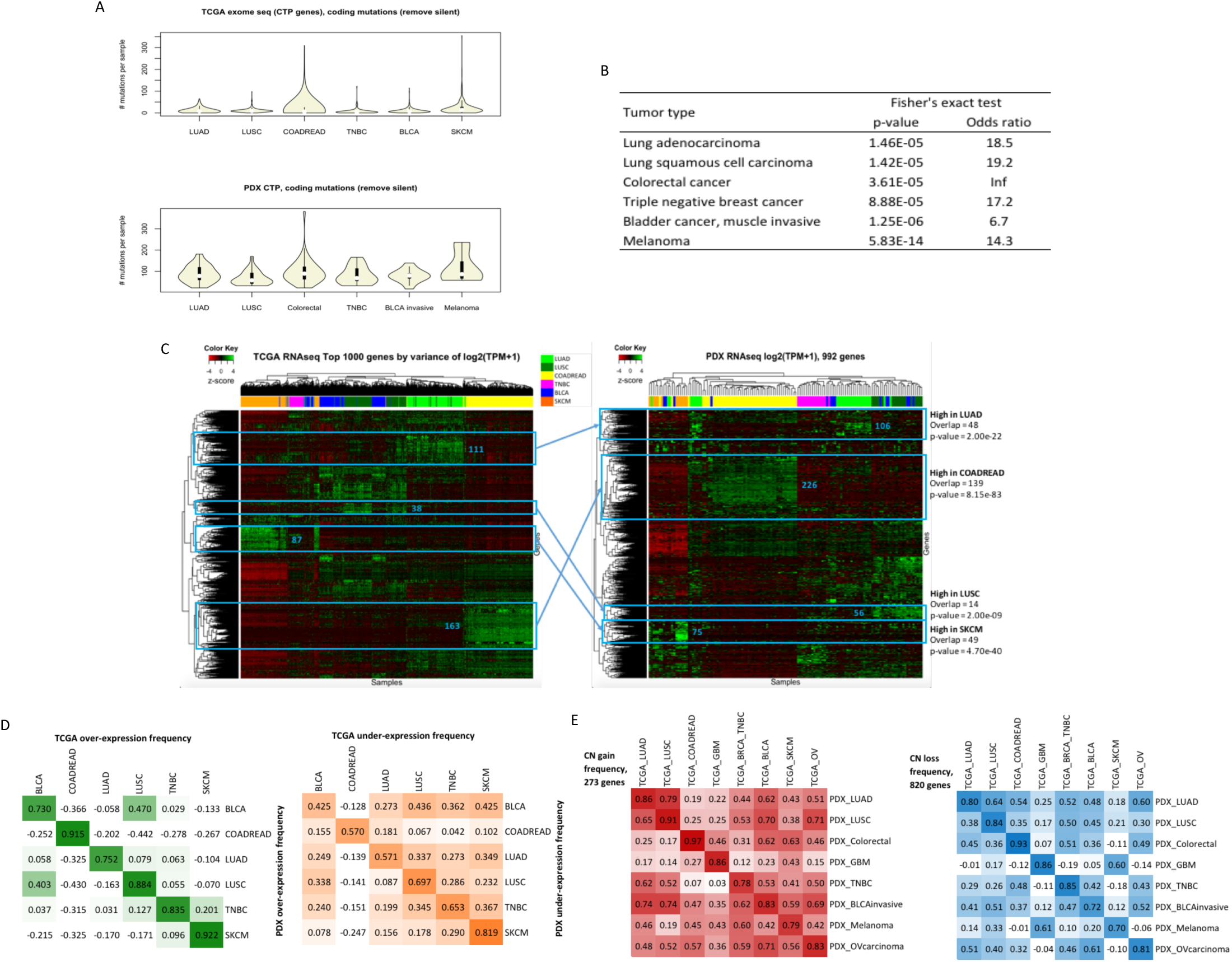
(A) Distribution of mutational load per sample of non-silent coding somatic mutations of CTP genes from exome sequencing of TCGA samples and from CTP-panel sequencing of PDX models (all filters included). (B) Overlap of CTP genes that have non-silent coding somatic mutations with >5% mutation frequency in TCGA data with genes that have at least one non-silent coding somatic mutation in PDX CTP data (all filters and rescue of clinically relevant variants included) for each tumor type. Fisher’s exact test is used to compute the significance of the overlap. (C) Hierarchical clustering of z-score of expression (log_2_(TPM+1)) of top 1000 most varying genes of TCGA RNA-Seq samples across different tumor types. The same set of genes (omitting non-expressed genes) is used to cluster the expression z-score by Hierarchical clustering of PDX RNA-Seq models across different tumor types. Gene sets were identified to be high expression in specific tumor types TCGA and PDX separately and were found to share significant overlap. (D) Correlation frequency of genes that are over-expressed (z-score of log_2_(TPM+1) > 1, green) or under-expressed (z-score of log_2_(TPM+1) < −1, orange) across each tumor type between PDX models and TCGA samples. (E) Correlation of frequency of copy number gain (red) or loss (blue) of selected genes frequently amplified or deleted in TCGA tumors predicted by GISTIC analysis for each tumor type between PDX and TCGA datasets.

#### Expression signatures of primary patient tumors in TCGA recapitulated in the PDX resource

The top 1000 most varying genes by expression z-scores in 6 TCGA tumor types (Supplementary Table S10) were able to independently cluster both TCGA samples and the PDX samples by their tumor types (Figure 6C). We observed clusters of genes that were highly expressed in specific tumor types in TCGA were recapitulated in the PDX expression data (hypergeometric p-value < 1×10^−8^), which demonstrated the replicability of TCGA expression signatures in the PDX resource. The frequencies of over- and under-expression for the top-varying genes for each tumor type displayed better correlation for the same tumor type for PDX versus TCGA compared to other tumor types (Figure 6D). The varying level of concordance between different tumor types in TCGA data was also maintained in the PDX versus TCGA comparison (Supplementary Figure S11). Alternatively, the differentially expressed genes of each tumor type versus all other tumors within the TCGA or PDX samples displayed significant overlaps (p<1×e^−6^), despite different sample sizes and different proportion of tumor types (Supplementary Table S11).

#### Copy number profiles of primary patient tumors in TCGA recapitulated in PDX resource

We showed that the frequency of genome-wide copy number aberrations for each tumor type in the PDX resource (Supplementary Table S12, Supplementary Figure S12) were similar to the primary tumors in TCGA (Supplementary Figure S13). Moreover, the PDX tumors had the highest correlation in gain and loss frequencies of significantly amplified and deleted genes for the same tumor type in TCGA compared to other tumor types (Figure 6E and Supplementary Figure S14A). The varying levels of correlation between different tumor types were preserved between the TCGA versus TCGA tumors and the TCGA versus PDX tumors (Figure 6E and Supplementary Figure S15B). Consistent with the earlier observations, there was a weaker concordance with TCGA data when amplification and deletion was called relative to the diploid state (Supplementary Figure S14B).

## Discussion

The application of PDX models in pre-clinical research and personalized therapy requires that the engrafted human tumors are accurately characterized for tumor-specific mutations [3]. The development of bioinformatics workflows to call somatic mutations (SNVs, Indels), copy number aberrations and gene expression from PDX sequencing or array data requires balancing sensitivity and specificity [22, 30], especially when paired normal samples for engrafted tumors are not available. Using genomic and transcriptomic data from models in the JAX PDX Resource, we conducted a systematic analysis to address several key data analysis challenges and tailored our workflows to optimize the sensitivity and specificity of the results.

Our recommendations for the somatic mutation calling from PDX DNA sequencing data in the absence of paired-normal samples are as follows:

- Remove mouse reads with Xenome (or equivalent) to eliminate variants called from mouse reads mapping to the human reference genome
- Filter with germline variant databases to improve somatic mutation calling
- Filter highly recurrent mutations to remove false positives arising from sequencing or analysis related errors
- Rescue clinically relevant variants which were filtered in the upstream steps as they were likely to be present as important mutations in the tumor

Despite implementing multiple filters to remove putative germline and other FP mutations, the mutation rate remains higher in PDX tumor types when compared to TCGA. One possible reason for this difference that is not related to the informatics challenges described in this paper is that many of the human tumor samples used to generate PDX models arose from metastatic lesions and from patients with prior treatment whereas many of the tumor samples used for TCGA were early stage tumors. PDX tumors were thus expected to harbor more mutations due to longer tumor evolution [36, 37]. Also, previous studies have noted that PDX engraftment success is better for late stage tumors that are likely to have more aggressive phenotypes than early stage tumors [38, 39]. As such, there is a likelihood for biased selection towards such tumor subtypes in the engrafted tumors that are known to harbor more mutations than tumors from early stages.

For evaluation of gene expression differences in individual tumors, matched normal tissue is ideal but not available for PDX models in the JAX PDX Resource. To compare gene expression among the engrafted tumors, we used expression z-scores across all tumor types as the best proxy for calling over- and under-expression. In a subset of PDX samples in which both expression and copy number data are available, we estimated the “normal” expression of each gene with the average expression for samples with normal copy number state, given that sufficient samples are available for the tumor type. While this approach neglects other mechanisms of gene regulation, we were able to better estimate the normal expression for some genes like MYC which tends to be frequently amplified and over-expressed across many tumor types. For copy number, we defined the “normal” state of each PDX tumor using the estimated ploidy to call relative gain and losses as this takes into account errors in ploidy estimation.

As one approach to assessing the results of our genomic characterization workflows, we compared the JAX PDX models with patient cohorts in TCGA at the genomic and transcriptomic level. Other than small differences in genomic mutations, the engrafted PDX tumors reflected the human tumors in copy number variations and gene expression. Using colorectal cancer as an example, we demonstrated that the integration of different data types showed that known perturbed pathways in cancer were altered in a consistent manner across PDX and TCGA tumors (Supplementary Figure S16), with similar combinations of alterations occuring at comparable frequencies. Taken together, we have created a set of workflows for the analysis of genomic and transcriptomic data from PDX tumors that have no paired normal sample to reliably identify true somatic mutations and expression changes.

## Acknowledgements

The workflows reported in this publication was partially supported by the National Cancer Institute of the National Institutes of Health under Award Number P30CA034196. The content is solely the responsibility of the authors and does not necessarily represent the official views of the National Institutes of Health. The public portal for JAX PDX data is supported by R01CA089713.

## Methods

### Genomic and transcriptomic profiling of samples

#### DNA sequencing

Flash frozen tissues were pulverized using a Bessman Tissue Pulverizer (Spectrum Chemical) and homogenized in Nuclei Lysis Buffer (Promega) using a gentleMACS dissociator (Miltenyi Biotec Inc). DNA was isolated using the Wizard Genomic DNA Purification Kit (Promega) according to manufacturer’s protocols. DNA quality and concentration were assessed using a Nanodrop 2000 spectrophotometer (Thermo Scientific), a Qubit dsDNA BR Assay Kit on a Qubit Fluorometer (Thermo Scientific), and the Genomic DNA ScreenTape on a 4200 TapeStation (Agilent Technologies). Libraries were prepared using the Hyper Prep Kit (KAPA Biosystems) and SureSelectXT Target Enrichment System with the JAX Cancer Treatment Profile (CTP) targeted panel (Agilent Technologies), according to the manufacturer’s instructions. Briefly, the protocol entails shearing the DNA using the Covaris E220 Focused-ultrasonicator (Covaris), ligating Illumina specific adapters, and PCR amplification. Amplified DNA libraries are then hybridized to the CTP probes, amplified using indexed primers, and checked for quality and concentration using the High Sensitivity D5000 ScreenTape (Agilent Technologies) and Qubit dsDNA HS Assay Kit (Thermo Scientific). Libraries were pooled and sequenced 150 bp paired-end on the NextSeq 500 (Illumina) using NextSeq v2 reagents (Illumina).

#### RNA sequencing

Tissues preserved in RNAlater were homogenized in TRIzol (ThermoFisher Scientific) using a gentleMACS dissociator (Miltenyi Biotec Inc). Total RNA was isolated using the miRNeasy Mini kit (Qiagen) according to manufacturer’s protocols, including the optional DNase digest step. RNA quality and concentration were assessed using the RNA 6000 Nano LabChip assay on the 2100 Bioanalyzer instrument and Nanodrop 2000 spectrophotometer (Thermo Scientific). Prior to 2016, non-stranded libraries were constructed using TruSeq RNA Library Prep Kit v2 (Illumina). Starting in 2016, stranded libraries were prepared by the Genome Technologies core facility at The Jackson Laboratory using the KAPA mRNA HyperPrep Kit (KAPA Biosystems), according to the manufacturer’s instructions. Briefly, the protocol entails isolation of polyA containing mRNA using oligo-dT magnetic beads, RNA fragmentation, first and second strand cDNA synthesis, ligation of Illumina-specific adapters containing a unique barcode sequence for each library, and PCR amplification. Libraries were checked for quality and concentration using the DNA 1000 assay (Agilent Technologies) and quantitative PCR (KAPA Biosystems), according to the manufacturers’ instructions. Libraries were pooled and sequenced 75 bp paired-end on the NextSeq 500 (Illumina) using NextSeq High Output Kit v2 reagents (Illumina), or 100 bp paired-end on the HiSeq2500 (Illumina) using TruSeq SBS v3 reagents (Illumina).

#### SNP array

DNA samples were sent to the Genotyping Core at the Hussman Institute for Human Genomics (University of Miami) for genotyping on the Genome-Wide Human SNP Array 6.0 (Affymetrix). Quality control on the CEL files was carried out using the standard Contrast QC metric from the Affymetrix Genome Wide SNP 6.0 array manual.

### Somatic point mutation and indel calling workflow

#### Preprocessing and removal of mouse reads

DNA sequence data generated from PDX tumors underwent initial data processing as follows: (i) sequence reads with 70% of the bases having a quality score <30 (Q30) were discarded, (ii) bases with quality scores less than Q30 were trimmed from the 3’ end of the read, (iii) sequence reads with <70% of bases remain after trimming were discarded, (iv) both reads from pair-end sequencing were discarded if either read was discarded. If <50% of the total reads remained following the preprocessing steps, the sample was removed from the analysis. Following the initial data processing step described above, mouse reads were identified and filtered out using Xenome v1.0.0 [13]. Only read pairs with both reads classified as human were included in further analyses.

Sequence reads that passed all pre-processing steps were mapped to the reference human genome (build GRCh38.p5 with 262 alternate loci) using the BWA-MEM alignment tool with ALT-Aware mapping (Supplementary Figure S14) [40, 41]. Because low sequence coverage leads to poor sensitivity in variant calling, samples with less than 75% of the target region covered at least at ≥100X by human reads were excluded from further analysis.

#### Variant calling

The GATK best practices workflow (https://gatkforums.broadinstitute.org/gatk/categories/best-practices-workflows) using the UnifiedGenotyper, was used for variant discovery analysis [42-44], which is comprised of the following steps: (i) sorting the SAM/BAM file by coordinate, (ii) removing duplicates to mitigate biases introduced by library preparation steps such as PCR amplification by Picard (https://broadinstitute.github.io/picard/), and (iii) recalibrating the base quality scores as the variant calling algorithms rely heavily on the quality scores assigned to the individual base calls in each sequence read. Pindel [32] was also incorporated into the workflow to call indels that have been missed by the GATK UnifiedGenotyper.

#### Quality filtering of variants for targeted sequencing

High quality variants from both variant callers in the PDX samples were obtained based on GATK hard filtering (see below), and have a read depth (DP) of ≥140 and allele frequency (ALT_AF) of ≥5%. These DP and ALT_AF thresholds were optimized using a set of known and validated mutations and samples reported earlier for the JAX CTP targeted panel sequencing at high coverage (average 941X) [45]. The parameters for GATK hard filtering [29] were set as default as recommended by GATK best practices (https://software.broadinstitute.org/gatk/documentation/article.php?id=6925, https://software.broadinstitute.org/gatk/documentation/article.php?id=3225, https://gatkforums.broadinstitute.org/gatk/discussion/2806/howto-apply-hard-filters-to-a-call-set):

i. for point mutations, QD < 2.0, FS > 60.0, MQ < 40.0, MQRankSum < −12.5,ReadPosRankSum < −8.0
ii. for indels, QD < 2.0, FS > 200.0, ReadPosRankSum < −20.0. In addition, we verified that these default thresholds were able to detect all the known mutations in the CTP samples [45]. The average number of variants before and after quality filtering across the CTP samples is shown Supplementary Table S4.

#### Annotation of variants

Variants were annotated for their effect (gene, consequence, amino acid change, etc.) using SnpEff v4.3 [46] based on gene annotations from Ensembl (version GRCh38.84) and information from COSMIC version 80 [47], dbSNP build 144 [48]. The observed variant allele frequency in 1000 Genomes Project [49] and ExAC version 0.3 [45, 50] database were obtained using SnpSift tool by utilizing dbNSFP3.2a.txt database. We further annotated each variant with 1) known or predicted gain or loss of protein function, 2) potential treatment approach for any cancer type and 3) drug sensitivity and resistance effects in clinical or preclinical studies, based on curated clinical information from the JAX clinical knowledge base (CKB, https://ckb.jax.org/) [31]. The average number of variants annotated to be clinically relevant across the CTP samples is shown in Supplementary Table S4.

#### Filtering of germline variants

Since normal samples were unavailable for patients whose tumors were used to generate the PDX models, we generated a dataset of putative human germline variants using data from several public resources: (i) dbSNP, (ii) 1000 Genomes Project, (iii) ExAC database with MAF ≥1%, and (iv) a compendium of variants from 20 normal blood samples that were prepped and sequenced on the CTP panel using the same protocol as the PDX samples, with a frequency of 2/20 in normal samples or 1/20 in normal samples and 2/20 in PDX models. The number of variants in each of these databases are shown in Supplementary Table S3. The variants identified via GATK and Pindel in the PDX model tumors were annotated as germline and filtered out of the model’s somatic mutation calls if they were present in our aggregated dataset of putative germline variants and had allele frequencies between 40% to 60% or more than 90%.

#### Filtering putative false positives

Variants not in our aggregated dataset of putative germline variants described above but occurred at a frequency of 25% or greater across all PDX models (n=236) were considered to be putative false positive (FP) mutations. The rationale for this data filtering step was based on our observation that the maximum recurrent frequency of somatic mutated base positions was 6% across a compendium of TCGA tumor samples (n=3576, 9 tumor types that were also represented in the PDX model). Thus, we would expect that any mutated loci recurring across PDX samples at significantly higher rates to likely be FP. Systematic technical errors in sequencing and/or mapping are possible explanations for the common recurrent non-somatic mutations identified PDX models.

#### Rescuing the false negative variants

An exception to the germline and false positives exclusion process was made for variants (from GATK or Pindel) that were annotated as clinically relevant in JAX CKB. We rescued any filtered variants that were curated into the proprietary JAX-Clinical Knowledgebase (CKB, https://ckb.jax.org/) [31] with 1) known or predicted gain or loss of protein function, 2) potential treatment approach for any cancer type and 3) drug sensitivity and resistance effects in clinical or preclinical studies.

### Benchmarking of PDX somatic mutation workflow

To benchmark the PDX somatic mutation workflow, a simulated dataset (45 samples) was generated that included sequenced reads that includes sequencing errors of an Illumina HiSeq were generated *in-silico* for different samples with, 1) varying sequencing coverage, 2) spiked-in mutations to the reference human sequence representative of different tumor types, and 3) different proportions of spiked-in mouse reads (Supplementary Table S1).

#### Generation of simulated sequence reads

SeqMaker was used to generate simulated sequencing data based on human genome assembly GRCh38 with varying sequencing depth, read length, duplication rate, sequencing error and base quality range [51]. Reference sequences were extracted from target region of the CTP panel. Sequence reads for 5 samples were simulated using predicted mutations from PDX models of different cancer types from the CTP dataset to represent different spectrum of mutations, with a range of allele frequency to mimic germline and somatic mutations. For each simulated sample, we generated three technical replicates at 500X, 1000X and 1500X coverage.

#### Addition of mouse reads

Mouse sequencing reads were added in different fractions to the human-specific simulated dataset to mimic mouse contamination observed in PDX models. The mouse reads were extracted from the sequencing data of mouse DNA isolated from fresh spleen tissue of NSG mice on the CTP. For each simulated human-specific sample, we added mouse reads in three proportions (10, 15 and 25% of the total coverage).

#### Calculate sensitivity and specificity of mutation results based on different workflow filters

To evaluate the effect of each filter used in our workflow, we modified the somatic mutation workflow by: (i) omitting Xenome to filter mouse reads, and (ii) mapping to the reference sequence using BWA-MEM. Each modified workflow was used to process each PDX simulated library and each set of results, with and without quality filters, was used to compute the lists of true positive, false positive, true negative and false negative variants. As such, we can calculate the range of sensitivities and specificities of the predicted variants for all the simulated PDX models. We compared the distributions of precision, recall and F1-score (2*(Recall*Precision)/(Recall+Precision)) for different variations of the variant calling workflow on the simulated datasets. Furthermore, we compared the predicted allele frequencies of the true positives of each sample with the input by correlation.

### RNA-Seq expression workflow

#### Data processing and expression estimation

Prior to alignment to the human transcriptome, sequences from PDX tumors were processed for sequence quality. Only sequences with base qualities ≥30 over 70 percent of read length were used in downstream analyses. Quality trimmed reads were then analyzed using the default parameters of Xenome v1.0.0 (k=25) [13] to separate human, mouse, and ambiguous sequences (i.e., sequences that cannot be reliably classified as mouse or human). Sequence reads that passed the quality and Xenome screening were aligned to a human transcriptome dataset (ENSEMBL version GRCh38.84) using Bowtie v2.2.0 [52, 53]. Only samples with at least 1 million human reads were retained for expression analysis. Gene expression estimates were determined using RSEM v1.2.19 [54] (*rsem-calculate-expression*) with default parameters. We further normalized the expression estimate (expected_count from RSEM) using upper quantile normalization of non-zero expected counts and scaling to 1000.

### Classifier for EBV-associated PDX lymphomas

A gene signature for differentiating EBV-associated lymphomas was derived from the most highly differentially expressed genes between 20 EBV-associated lymphomas and 100 non-EBV tumors based on upper-quantile normalized RNA-Seq counts (RSEM). Gene set analysis on the resulting expression vector was performed with GSEA using the GenePattern webserver and default parameters (data not shown). 24 up-regulated and 24 downregulated genes from the set of differentially expressed genes were used to define the list of classifier genes (Supplementary Table S10). For each PDX sample, the upper-quantile normalized counts from RSEM of the classifier genes were transformed into z-scores using the mean and standard deviation computed across all PDX samples for each gene. Subsequently, a sign corresponding to the direction of regulation in the classifier table was multiplied to each z-score and the sum of these modified z-scores resulted in a single score for each PDX sample. A classifier score of >3.0 was used to identify a PDX tumor sample as a potential EBV-associated lymphoma.

### Copy Number Variant (CNV) workflow

#### Assessing the effects of mouse DNA on SNP array

DNA of the NSG mouse was hybridized on the Affymetrix SNP 6.0 array, and the signal intensity was extracted from the CEL files using Affymetrix Power Tools (apt-cel-extract). The mouse content for each PDX sample was estimated by the mouse reads proportion computed by Xenome of the mutation calling pipeline for the CTP sequencing of the same PDX sample.

#### Single-tumor CNV analysis

PennCNV-Affy and Affymetrix Power Tools [55-57] were used to extract the B-allele frequency (BAF) and Log R Ratio (LRR) from the resulting CEL files of the Affymetrix Human SNP 6.0 array. Due to the absence of paired-normal samples, the allele-specific signal intensity for each PDX tumor were normalized relative to 300 randomly selected sex-matched Affymetrix Human SNP 6.0 array samples obtained from the International HapMap project [58]. The single tumor version of ASCAT 2.4.3 [59] was then used for GC correction, predictions of the heterozygous germline SNPs and estimation of ploidy, tumor content and copy number segments with allele-specific copy number.

#### Annotation of CNV segments

The resultant copy number segments were annotated with loss of heterozygosity (LOH) and log_2_ ratio of total copy number relative to diploid state (copy number 2) and predicted ploidy from ASCAT. A segment was defined as LOH when the major-allele copy number was ≥ 0.5 and the minor-allele copy number was ≤ 0.1. Gene-level copy number and LOH were estimated by intersecting the genome coordinates of copy number segments with genome coordinates of genes (Ensembl annotation version 84 for genome assembly GRCh38). In cases where a segment boundary was contained within a gene’s coordinates, the most conservative (lowest) estimate of copy number was used and the gene was annotated with the number of overlapping segments.

#### Defining copy number gain and loss

The low-level copy number gain or loss of a gene was defined by the log_2_ ratio of the copy number relative to the average ploidy of the sample or diploid state with a threshold of ±0.4 respectively. We compiled a list of genes with focal copy number aberrations that were significantly amplified (n=273) or deleted (n=820) in the 8 tumor types (Supplementary Table S8) from the GISTIC 2.0 analysis from the TCGA FireBrowse website (http://firebrowse.org/). Using this set of genes, we compared the proportion of genes that would be classified as gain and loss when using different baselines (diploid state 2 or ASCAT predicted ploidy) for PDX models listed in Supplementary Table S12.

#### Comparison of copy number aberrations with gene expression

Using annotations from the Cancer Census resource [33] we analyzed the relationship between copy number aberrations and gene expression using a list of 23 oncogenes that are commonly amplified in cancers and a list of 40 tumor suppressor genes that are commonly deleted in cancers. These genes were classified into copy number states of high-level loss (log2(CN/ploidy) < −1), normal (−1 ≤ log_2_(CN/ploidy) ≤ +1) and high-level gain (log_2_(CN/ploidy) > +1). The expression fold change of each gene was calculated as the log_2_(TPM+1) relative to the mean expression across PDX samples with a stringent normal copy number state (−0.4 ≤ log2(CN/ploidy) ≤ 0.4). The significance of expression changes of each gene for the entire PDX resource with copy number gain or loss relative to the normal state was calculated using the Student’s t-Test.

### Comparison between PDX and TCGA data

#### Somatic mutations

We calculated the distribution of mutational load (number of non-silent, coding mutations in exonic regions per sample) of the CTP genes for 6 tumor types with at least 10 models in the PDX resource (colorectal cancer, lung adenocarcinoma, lung squamous cell carcinoma, melanoma, bladder carcinoma and triple-negative breast cancer, Supplementary Table S5). MAF files for somatic mutations based on whole-exome sequencing of the TCGA samples of 6 tumor types [60-64] were obtained from TCGA Data Portal and were used to compute the mutation frequency for CTP genes only. The Fisher’s exact test was used to test the significance of overlap of mutated genes between the PDX resource and TCGA patient cohorts for each tumor type. The genes in each PDX resource were considered if they were mutated in at least one sample, while the genes in each TCGA tumor cohort were considered if they were mutated with at least 5% frequency, due to a much larger sample size.

#### RNA-Seq gene expression

6 tumor types with at least 10 models in the PDX resource were selected for comparison with TCGA (colorectal cancer, lung adenocarcinoma, lung squamous cell carcinoma, melanoma, bladder carcinoma and triple-negative breast cancer, Supplementary Table S10). The scaled estimate (TPM × 10^−6^) from the RNA-Seq data of 6 tumor types in TCGA [60-65] were obtained from the TCGA FireBrowse website (http://firebrowse.org/). Non-expressed genes across all tumor types were removed (log_2_(TPM+1) < 2), and the top 1000 most varying genes based on their z-scores of log_2_(TPM+1) across all tumor types were selected to cluster the samples by hierarchical clustering. The frequencies of over-expression and under-expression of each gene is defined by the z-scores of log_2_(TPM+1) of ±1. Correlation of the gene expression frequencies in each tumor type was computed using Pearson correlation. The differential gene expression of each tumor type compared to all other tumor types was computed using limma [66] based on log_2_(TPM+1) values. Up-regulated (adjusted p-value < 0.05, log (fold change of TPM+1) > 1 by limma) or down-regulated (adjusted p-value < 0.05, log (fold change of TPM+1) < −1 by limma) genes were obtained for the PDX resource and TCGA patient cohorts separately. The significance of overlap of each set of genes between PDX and TCGA RNA-Seq data was determined using hypergeometric p-value.

#### Copy number aberrations

8 tumor types with at least 10 models in the PDX resource (colorectal cancer, lung adenocarcinoma, lung squamous cell carcinoma, melanoma, glioblastoma multiforme, bladder carcinoma, triple-negative breast cancer and ovarian carcinoma, Supplementary Table S12) selected to compare with corresponding primary tumors in the TCGA [60-65, 67-69]. For PDX samples, the low-level copy number gain or loss of a gene was defined by the log_2_ ratio of the copy number relative to the average ploidy of the sample (or copy number state 2) with a threshold of ±0.4 respectively. The amplification or deletion calls of each gene for the TCGA samples were provided (loss=-1, normal=0, gain=1) by FireBrowse (http://firebrowse.org/). Using the list of genes with focal copy number aberrations that were significantly amplified (n=273) or deleted (n=820) in the 8 tumor types from the GISTIC 2.0 analysis from the TCGA FireBrowse website, we calculated the copy number gain and loss frequencies of these genes for each tumor type in the PDX resource and TCGA cohorts using the respective gain and loss calls.

---

**Figure S1.**
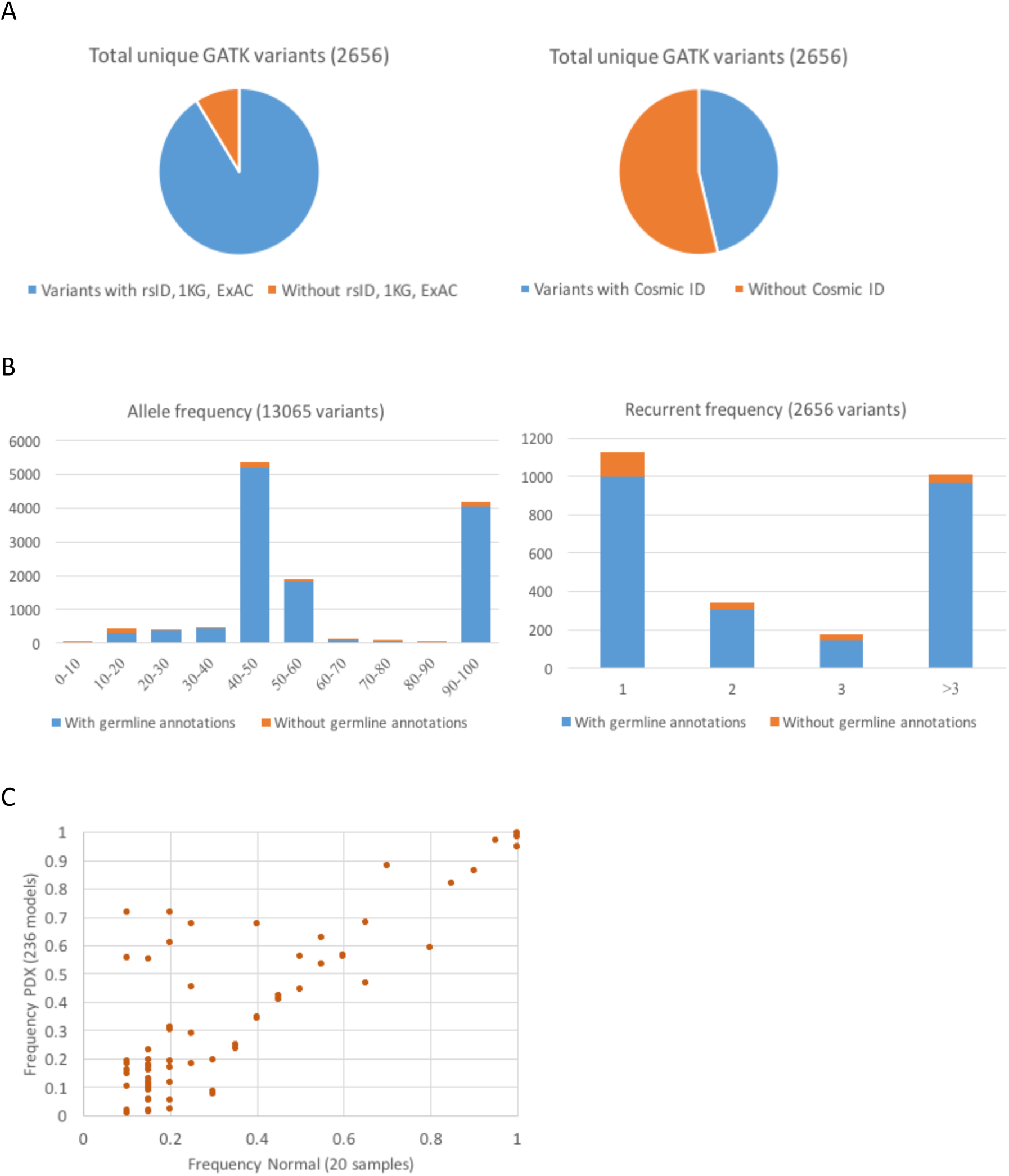
(A) This figure shows the annotation of variants of the 20 normal samples in JAX using public databases (dbSNP Build 144, 1000 Genomes, ExAC version 0.3, and COSMIC version 80) (B) The allele frequencies and recurrent frequencies of the variants. (C) Recurrent frequency of variants (> 1 sample) found in 20 normal samples and the corresponding recurrent frequency across 236 PDX models of different tumor types.

**Figure S2.**
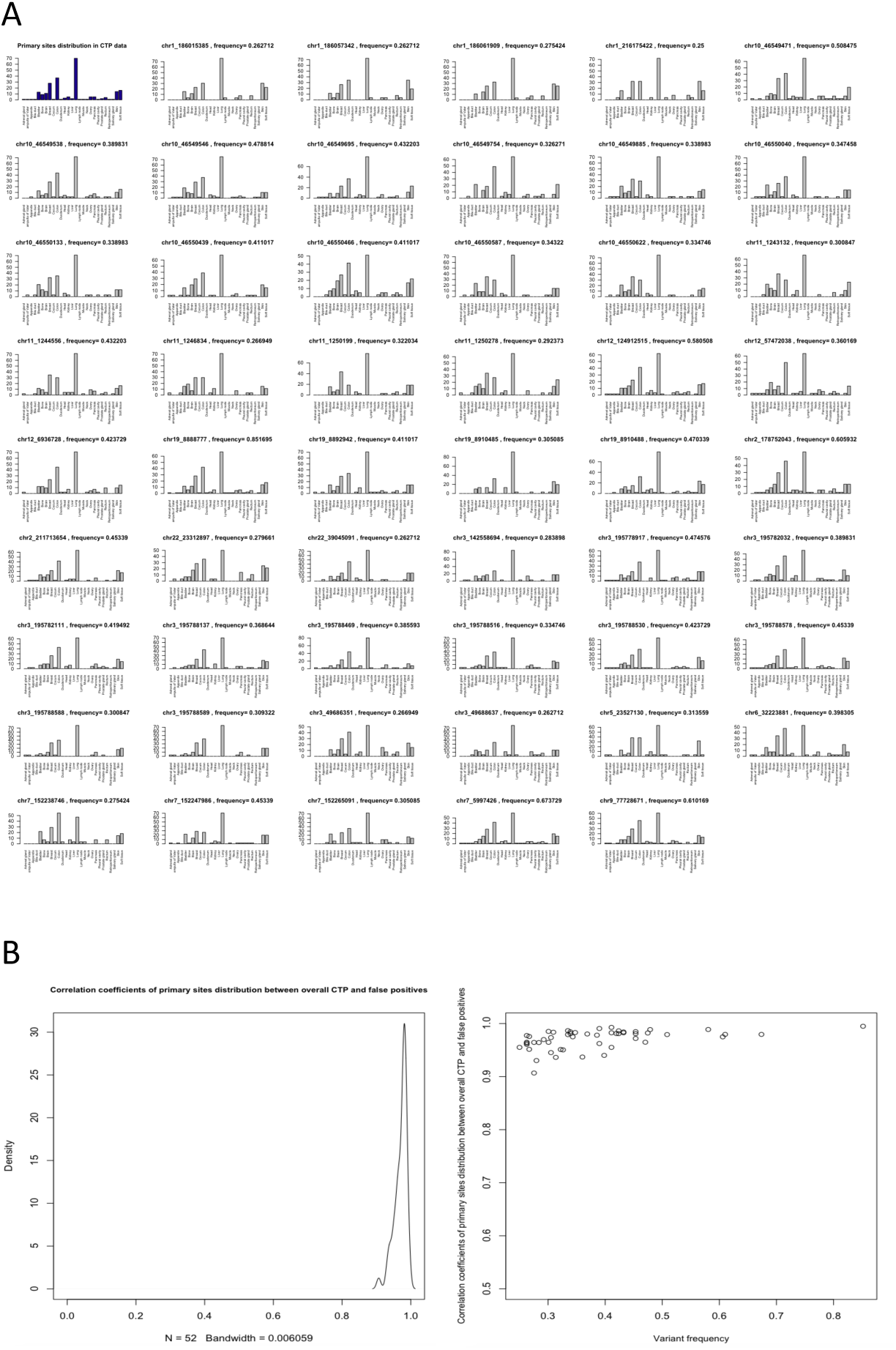
(A) Top left (dark blue): Distribution of tumor types across 236 PDX models; Others (grey): Frequency of tumor types for each frequently mutated position normalized by the recurrent frequency of the mutation. (B) Correlations of tumor type frequency for the 236 PDX models (dark blue in B) with the tumor type frequency for each frequently mutated position (grey in B).

**Figure S3.**
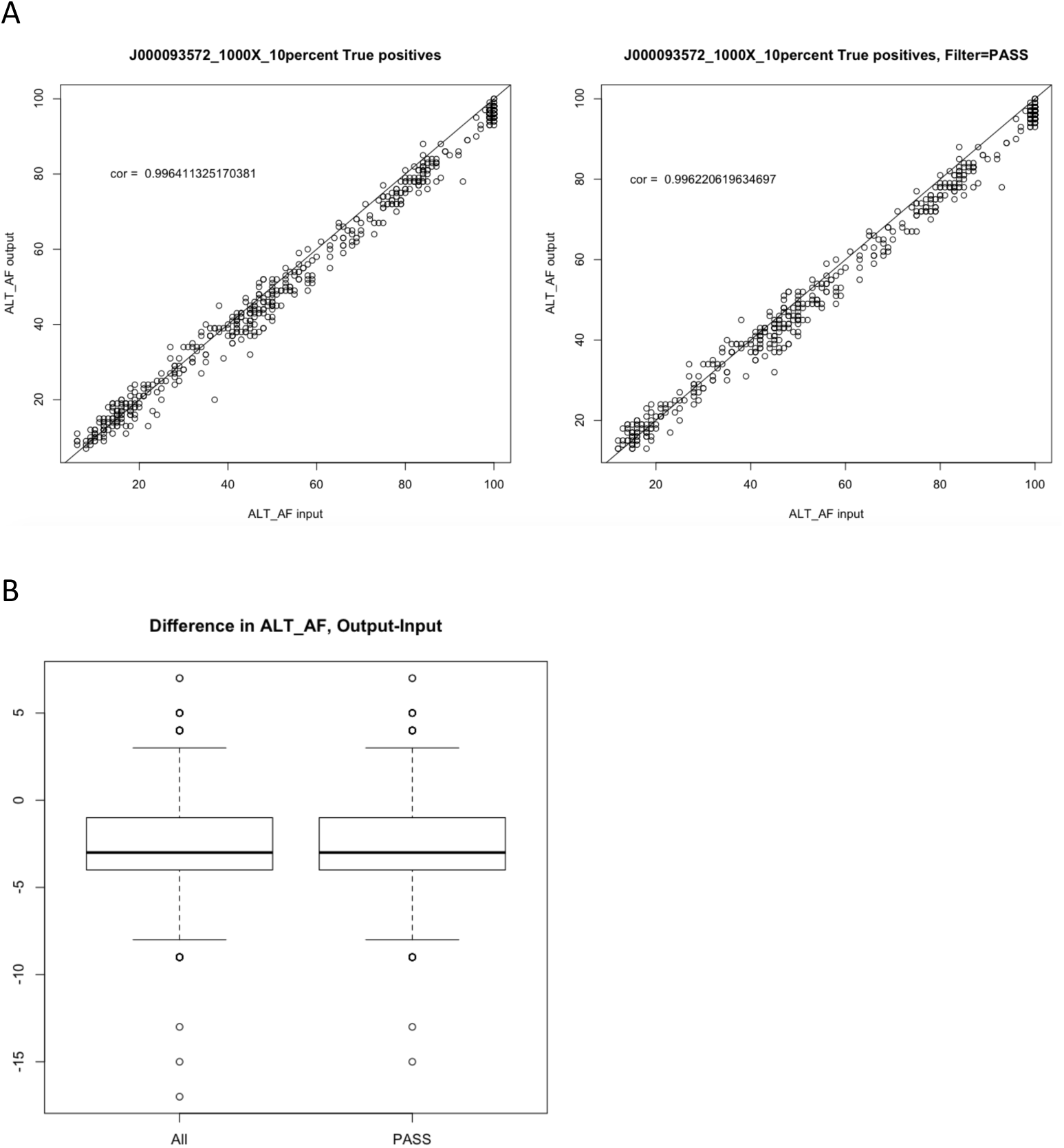
(A) Correlation of alternate allele frequencies between input and true positive variants for one of the simulated samples for the complete feature pipeline. ALL: all variants called by the pipeline; PASS: variants annotated as “PASS” in the pipeline which pass the hard filters, minimum read depth and minimum alternate allele frequency of the variant. The correlation coefficient for all simulated samples are found in Supplementary Table S3. (B) Difference in alternate allele frequencies between input and true positive variants for one of the simulated samples (J000093572_1000X_10percent) for the complete feature pipeline. ALL: all variants called by the pipeline; PASS: variants annotated as “PASS” in the pipeline which pass the hard filters, minimum read depth and minimum alternate allele frequency of the variant.

**Figure S4.**
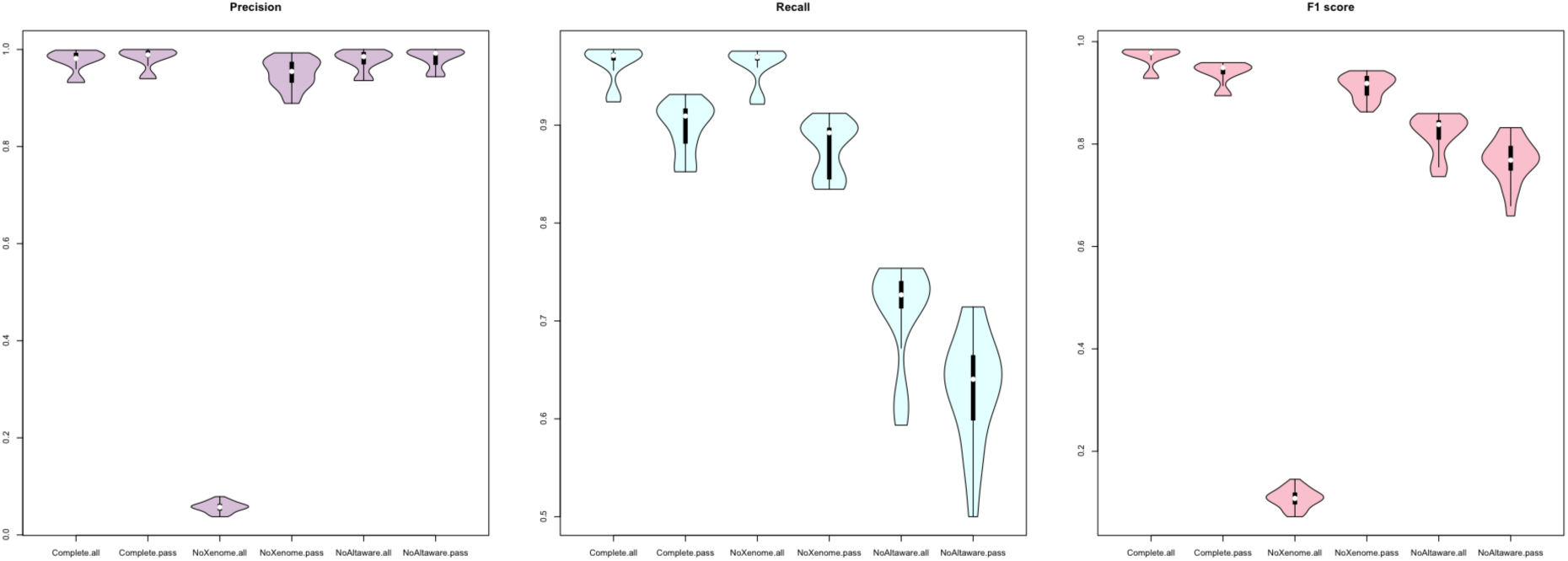
This figure shows the benchmarking of the CTP variant calling pipeline using 45 simulated sequencing datasets different samples, sequencing coverages, and mouse DNA content (see Supplementary Table S2) using precision, recall and F1 score based on the input variants for each sample. Complete: variant calling pipeline with all steps included; NoXenome: variant calling pipeline with Xenome omitted; NoAltaware: variant calling pipeline using hg38 reference with alternate sequences but using standard BWA for mapping instead of BWA-ALT-Aware; all: all variants called by the pipeline; pass: variants annotated as “PASS” in the pipeline which pass the hard filters, minimum read depth and minimum alternate allele frequency of the variant.

## Presence of alternate loci in the genome assembly

The GRCh38.p5 human genome assembly includes 262 regions of alternate loci to account for human chromosomal regions that exhibit sufficient variability to prevent adequate representation by a single sequence [29]. As such, we aligned the reads to both primary and alternate chromosomal reference sequences using BWA-MEM with ALT-aware. When alignment is performed using BWA-MEM only, the recall of the variants is much lower (∼30%) than the standard pipeline with or without hard-filtering (Supplementary Figure S14 and Supplementary Table S2). This shows that using an alignment tool not catered for alternate loci mapping reduces the overall sensitivity of the variant calling due to lesser reads being correctly mapped. The correlation of allele frequencies also decreases and the reduction in median allele frequency increases up to 15% (Supplementary Table S3).

**Figure S5.**
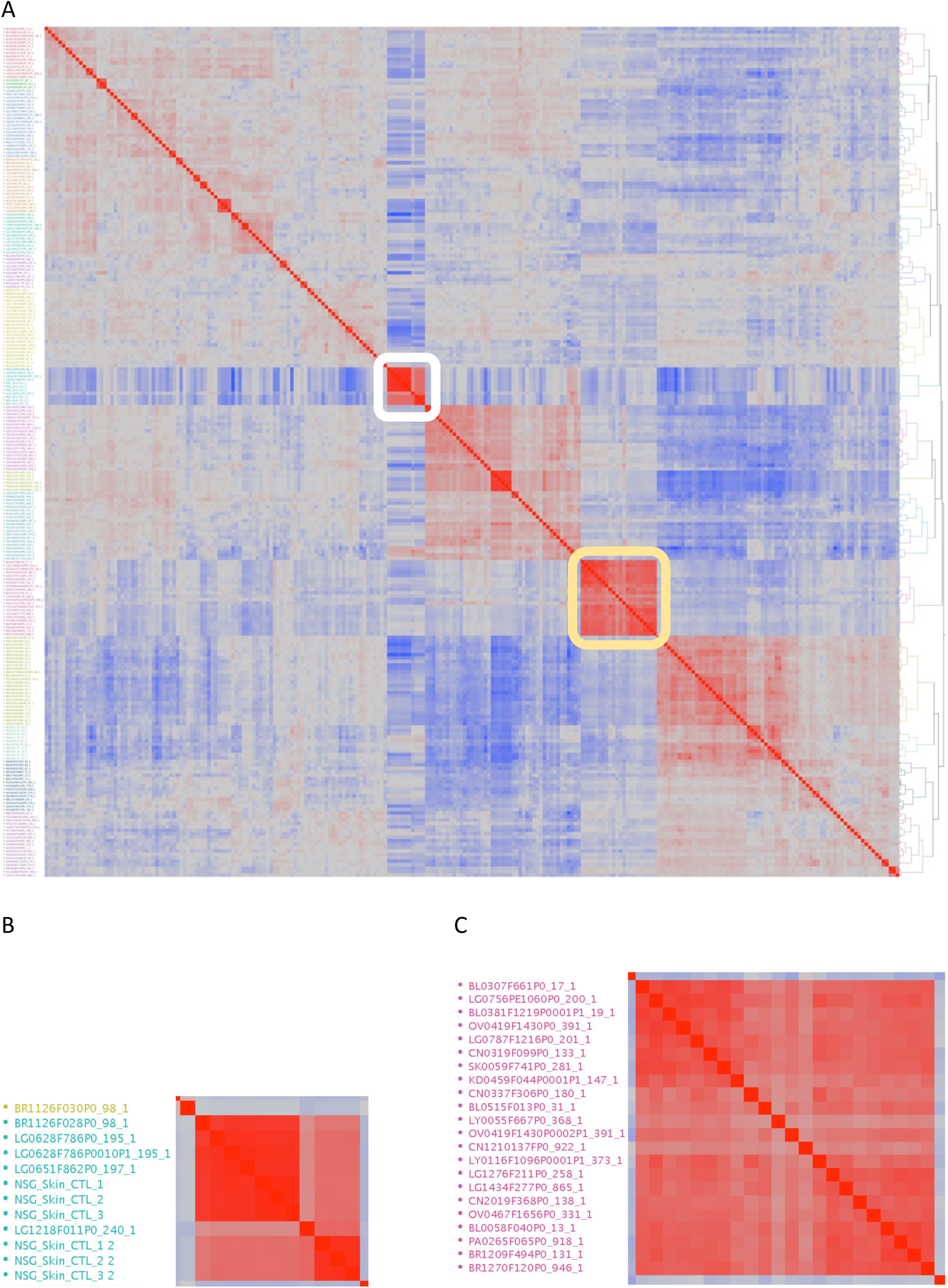
(A) Hierarchical clustering of the pairwise correlations of microarray expression between pairs of models, with red representing perfect correlation (+1), and blue perfect anti-correlation (−1). The two largest red blocks (highlighted in white and yellow) show the mouse introgressed and EBV transformed models. The other blocks, which much lower average correlation, typically show related tumor types (e.g., the lower right block is all neurological tumors). (B) A small fraction of tumors, highlighted in white in (A), that were heavily introgressed by mouse tissues were clustered with expression of NSG mouse (skin sample). (C) EBV Lymphoma models, highlight in yellow in (A), show an extremely highly correlated expression pattern regardless of the original tissue or tumor type.

**Figure S6.**
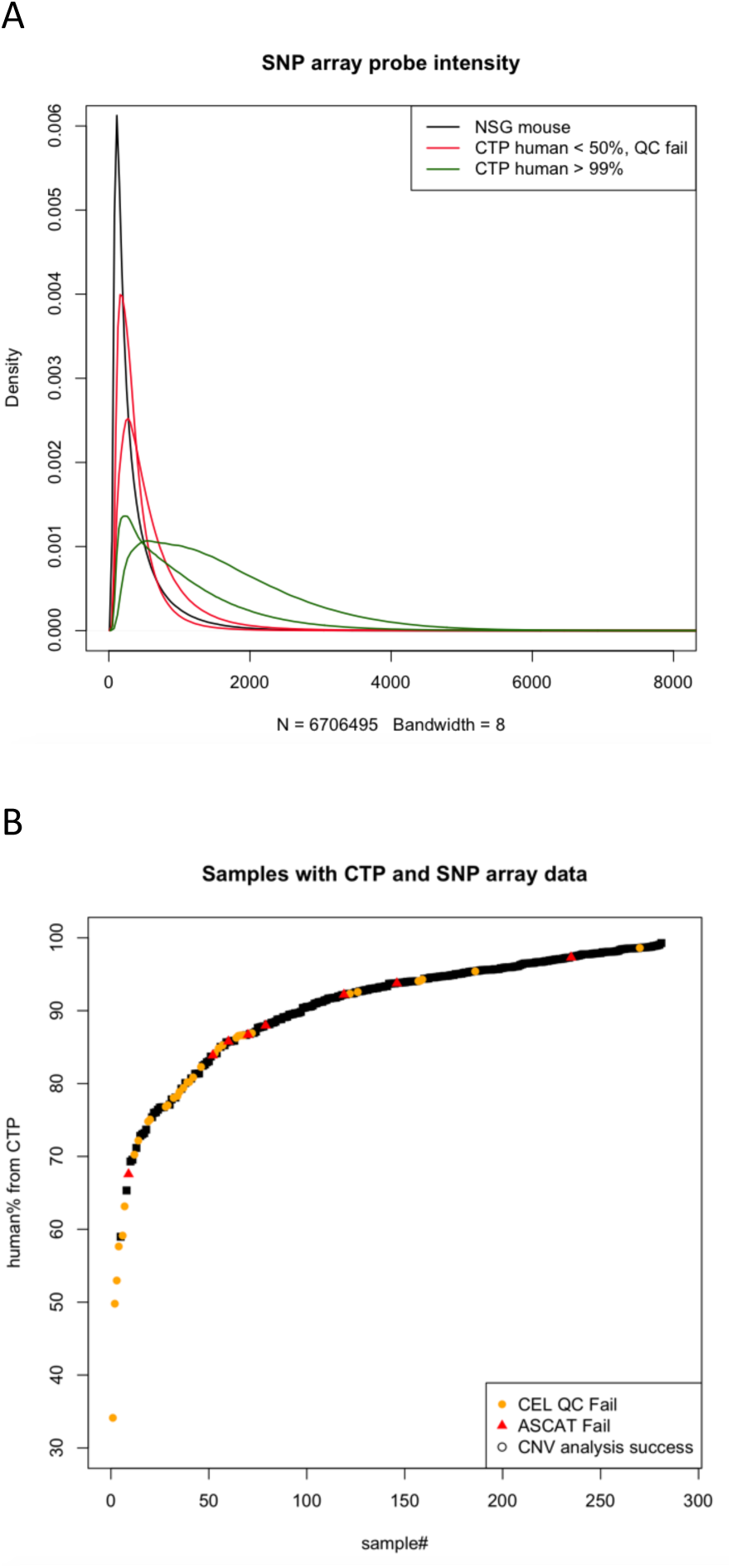
(A) Distribution of probe intensity of the SNP array for PDX samples with different mouse DNA content (using the percentage of human reads estimated from the CTP sequencing as a proxy for human DNA content on the SNP array): > 99% human DNA (green), < 50% human DNA with QC failure of SNP array CEL file (red), and 100% NSG mouse DNA (black). (B) Human DNA content of PDX samples classified by successful CNV prediction (black squares), failure in QC of CEL files (orange circles), and failure in ASCAT analysis red triangles).

**Figure S7.**
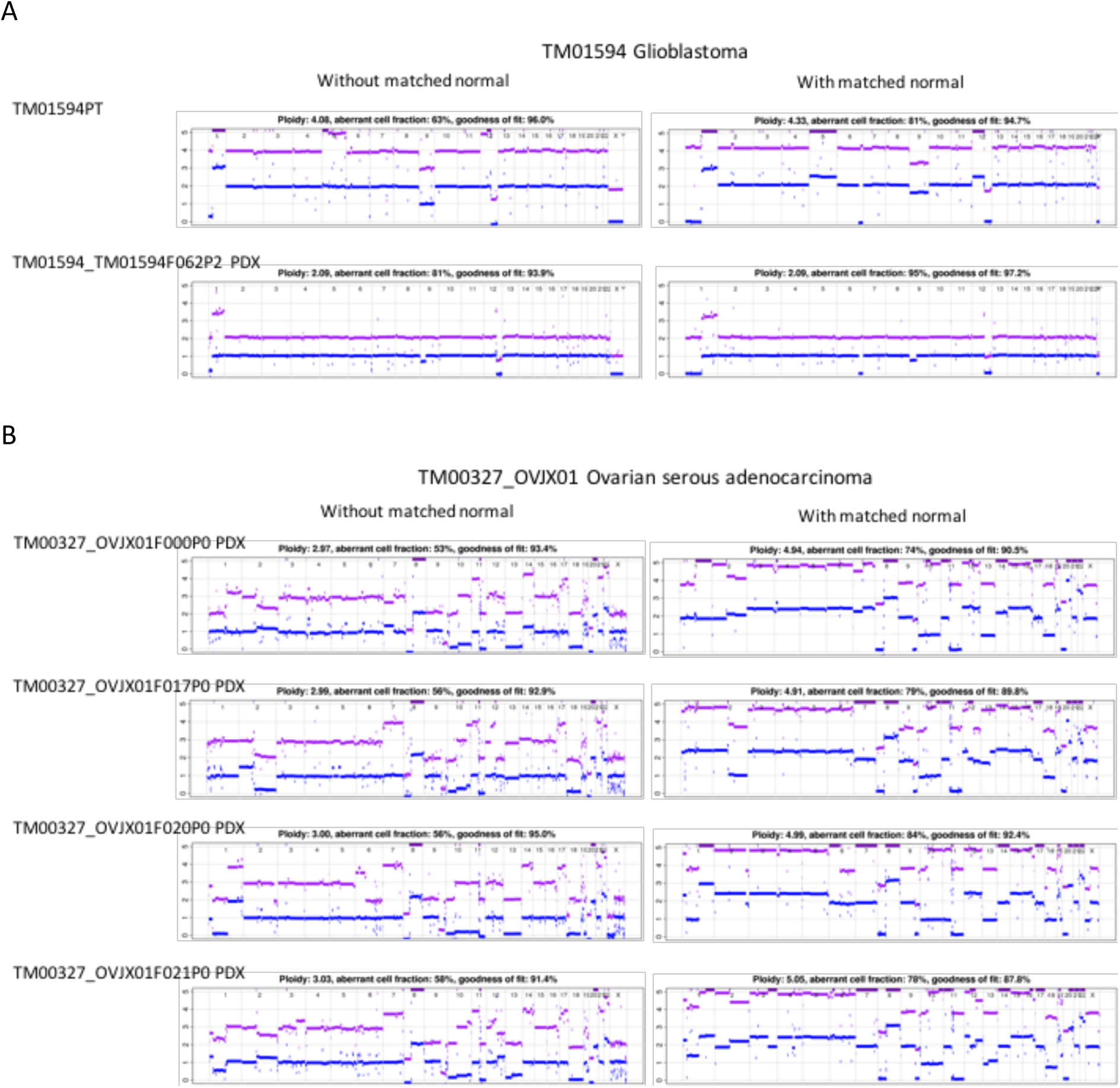
(A) And (B): CNV profiles of PDX models with matched models and multiple samples from the corresponding patient tumor, multiple passages or multiple samples (mouse) of same passage.

**Figure S8.**
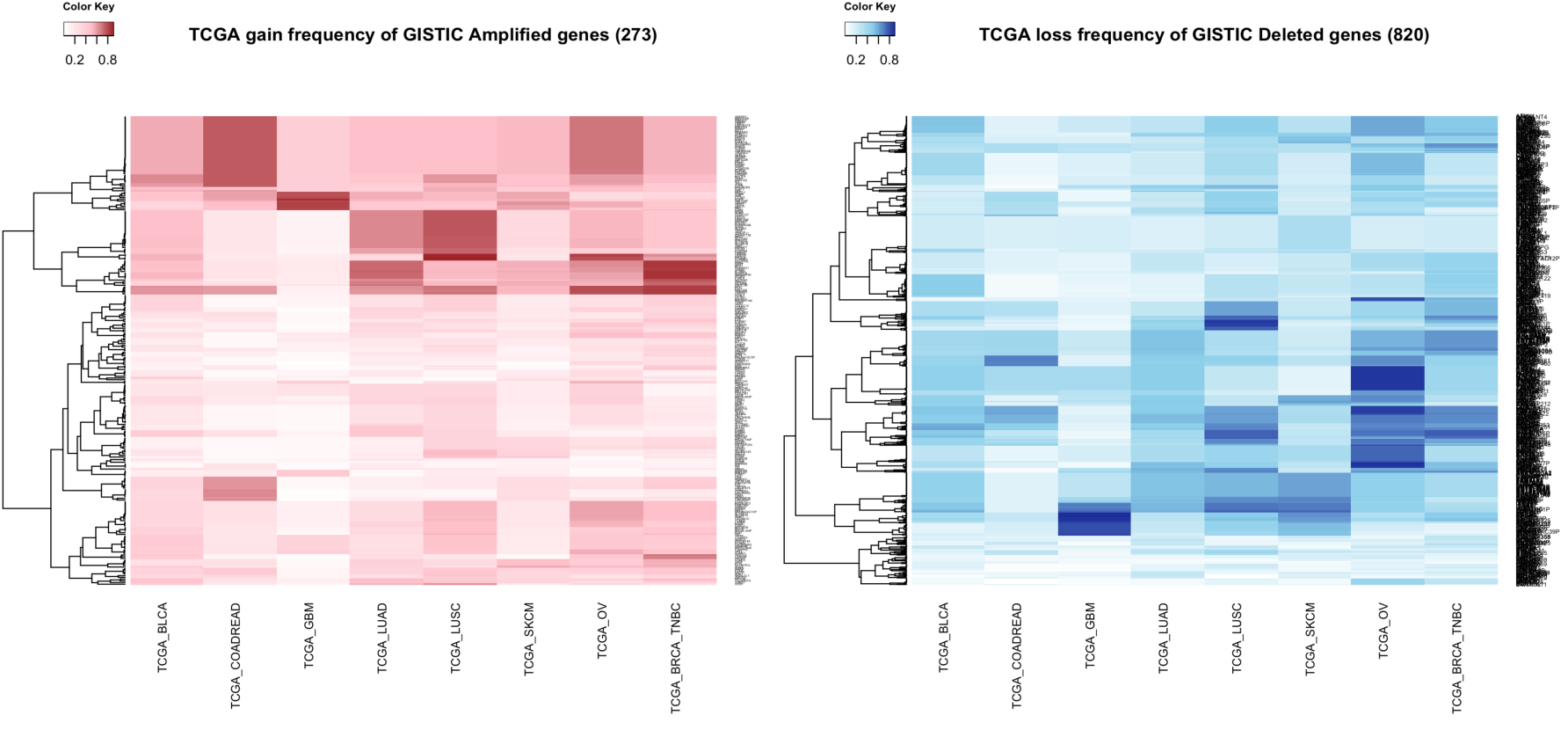
Frequency of copy number gain (red) or loss (blue) of selected genes frequently amplified or deleted in TCGA tumors predicted by GISTIC analysis for each tumor type in TCGA SNP array datasets.

**Figure S9.**
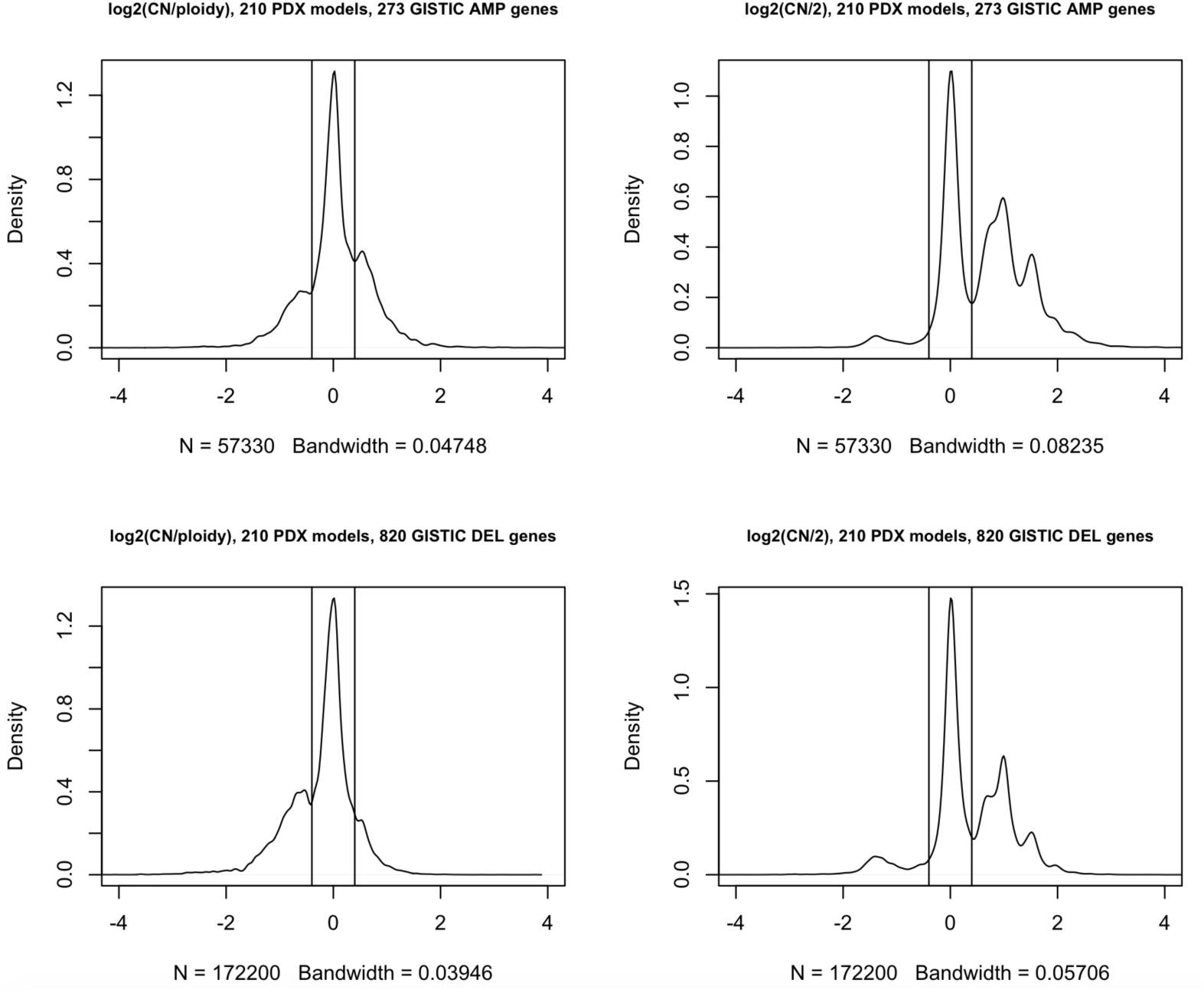
Distribution of log_2_ ratio of gene copy number relative to the estimated overall ploidy of each individual PDX sample or the diploid state for selected frequently amplified or deleted genes in TCGA tumors predicted by GISTIC analysis across all PDX samples. The threshold of low-level gain and loss is defined as log_2_(CN/ploidy) > +0.4 and log_2_(CN/ploidy) < −0.4 respectively.

**Figure S10.**
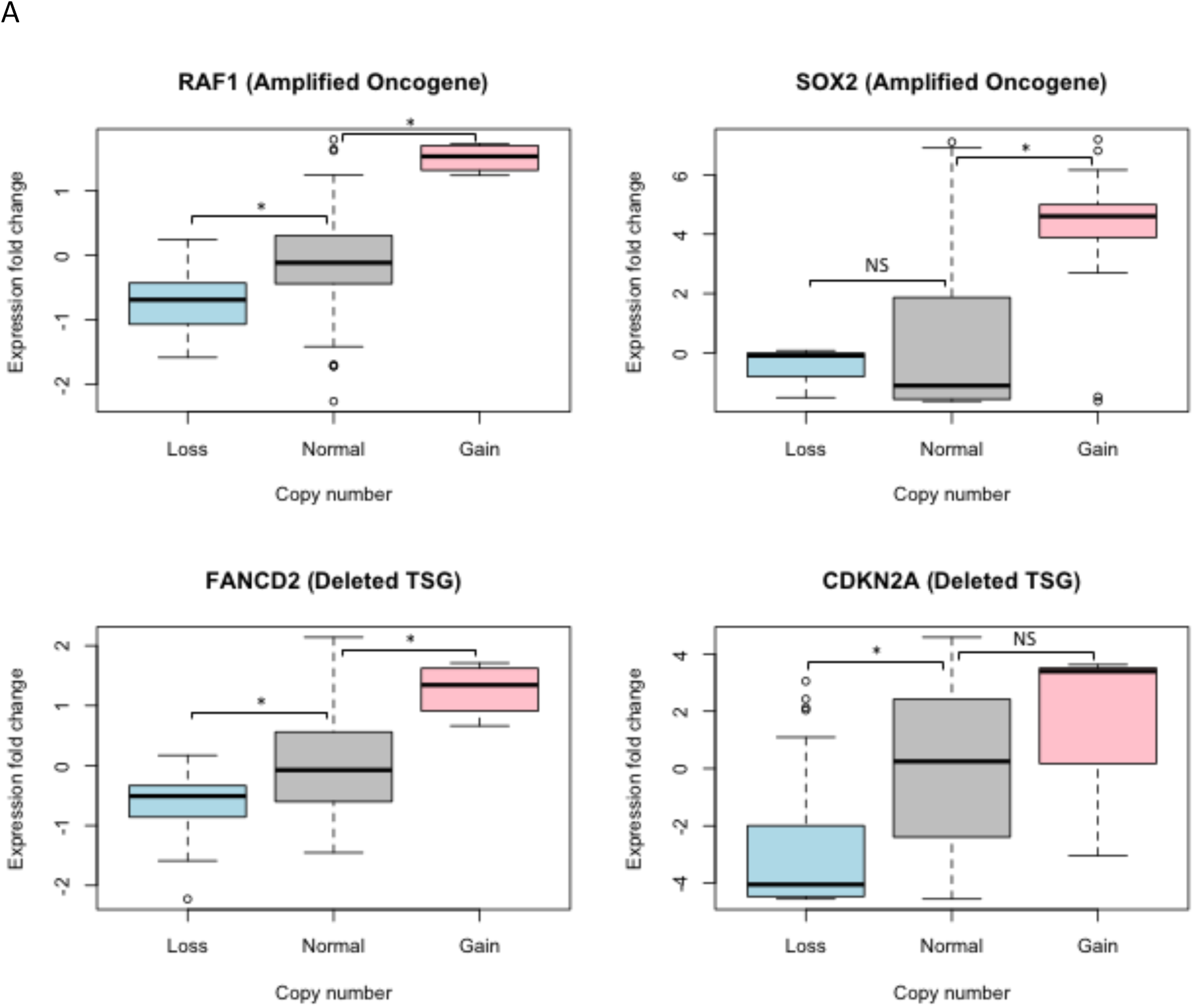
Expression fold change of each gene across all PDX samples, defined by fold change of log_2_(TPM+1) relative to the mean expression of samples with a stringent normal copy number state (−0.4 < log2(CN/ploidy) < 0.4). Here, a higher-level copy number gain and loss is defined as log_2_(CN/ploidy) > +1 and log_2_(CN/ploidy) < −1 respectively. The normal copy number state is defined as −1 < log_2_(CN/ploidy) < +1. Significance in differences in expression by Student’s t-test (*: p-value < 0.005, NS: non-significant).

**Figure S11.**
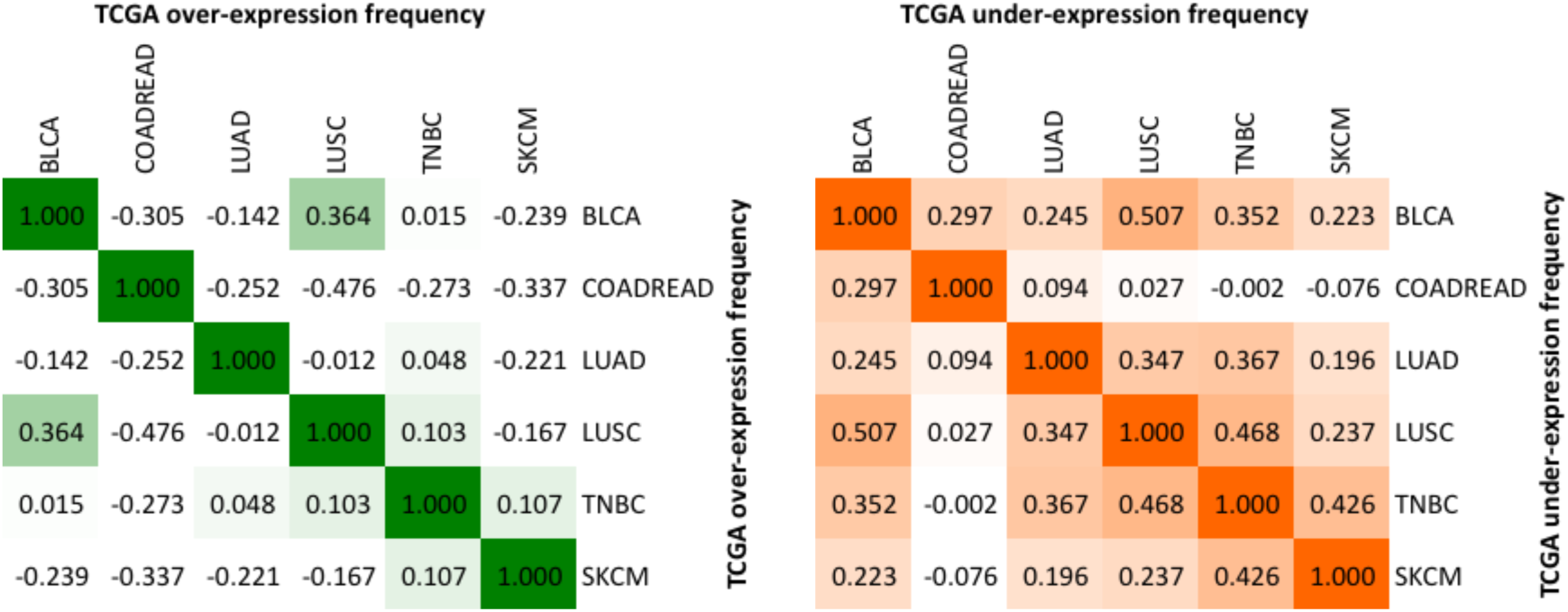
Correlation frequency of genes that are over-expressed (z-score of log_2_(TPM+1) > 1, green) or under-expressed (z-score of log_2_(TPM+1) < −1, orange) between each tumor type in TCGA RNA-Seq samples.

**Figure S12.**
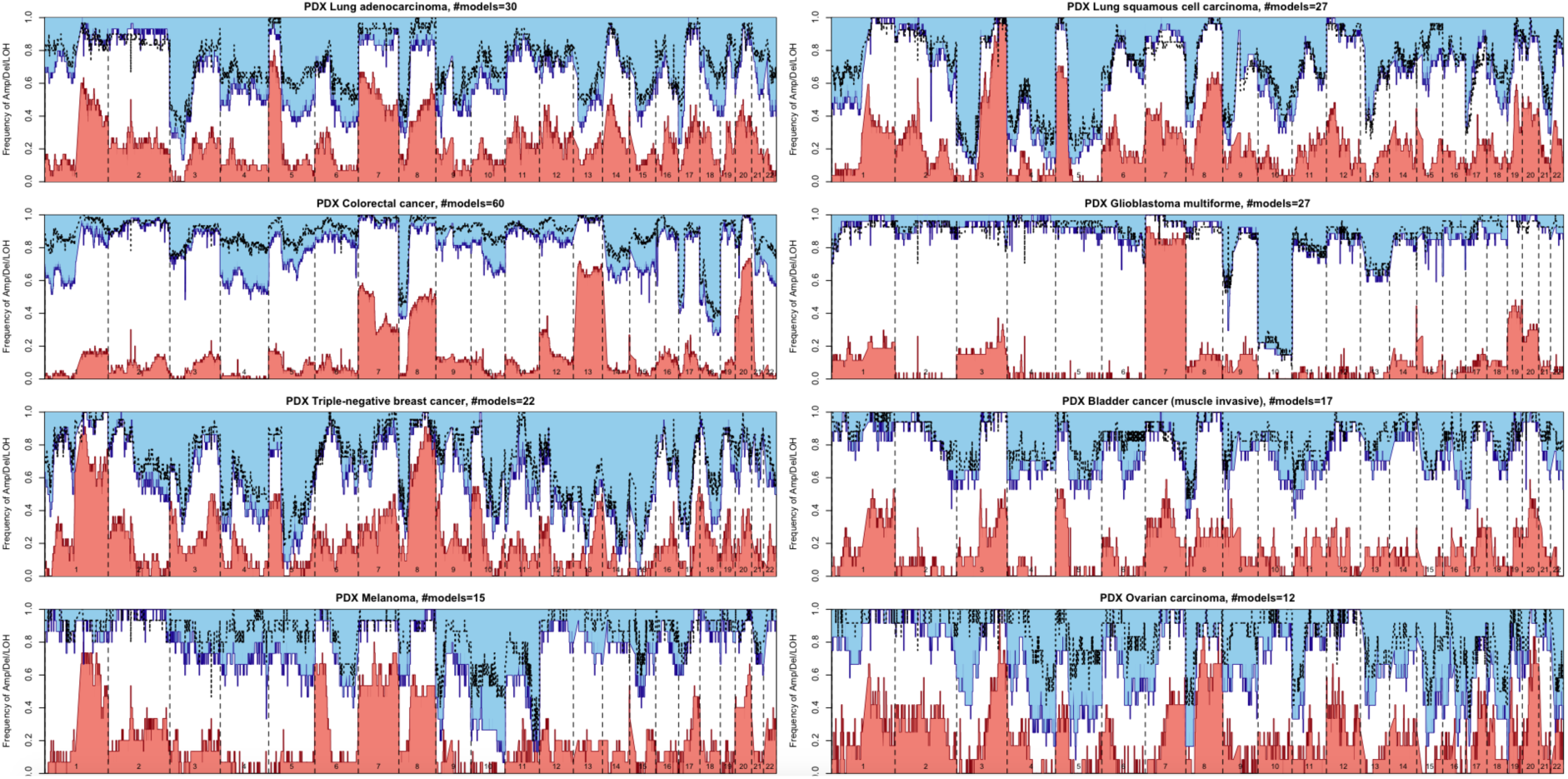
Frequency of genome-wide copy number gain, loss and LOH across PDX models for 8 tumor types.

**Figure S13.**
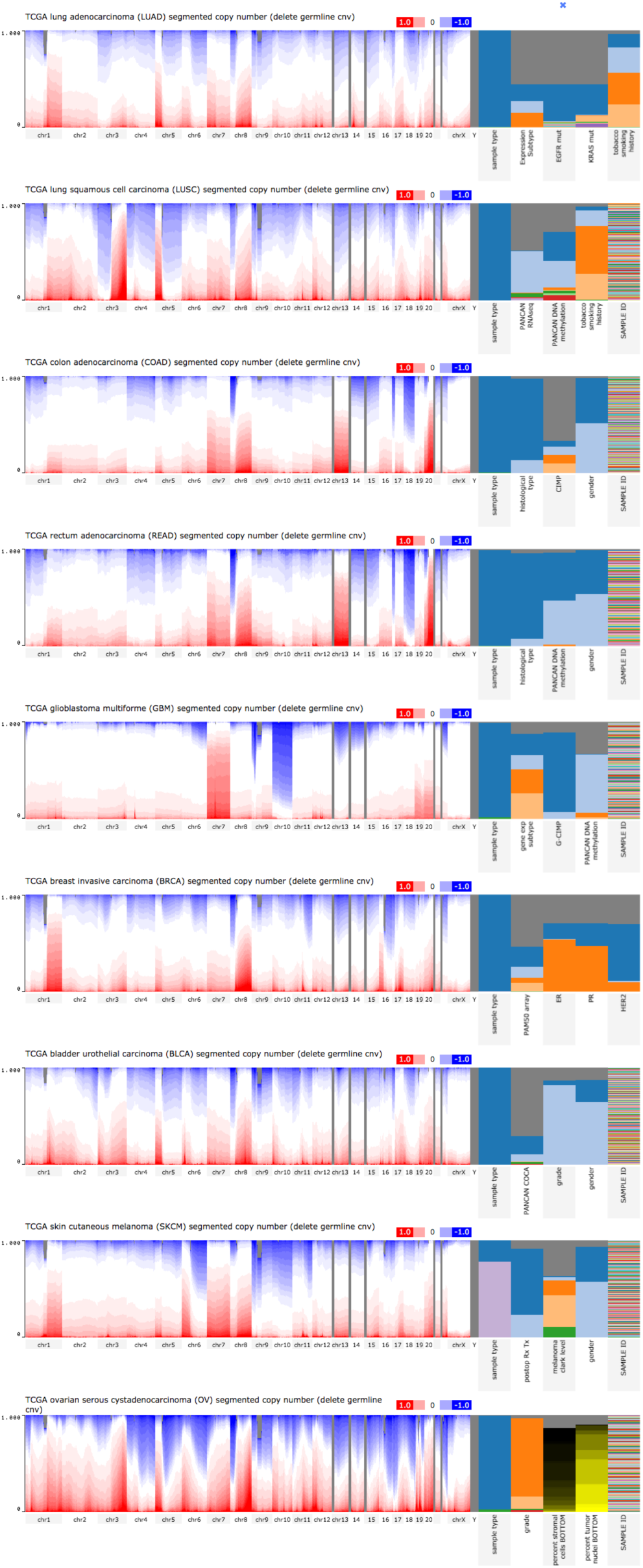
Frequency of genome-wide copy number gain and loss across TCGA samples for 8 tumor types. (Compiled from UCSC Cancer Browser, https://genome-cancer.ucsc.edu/)

**Figure S14.**
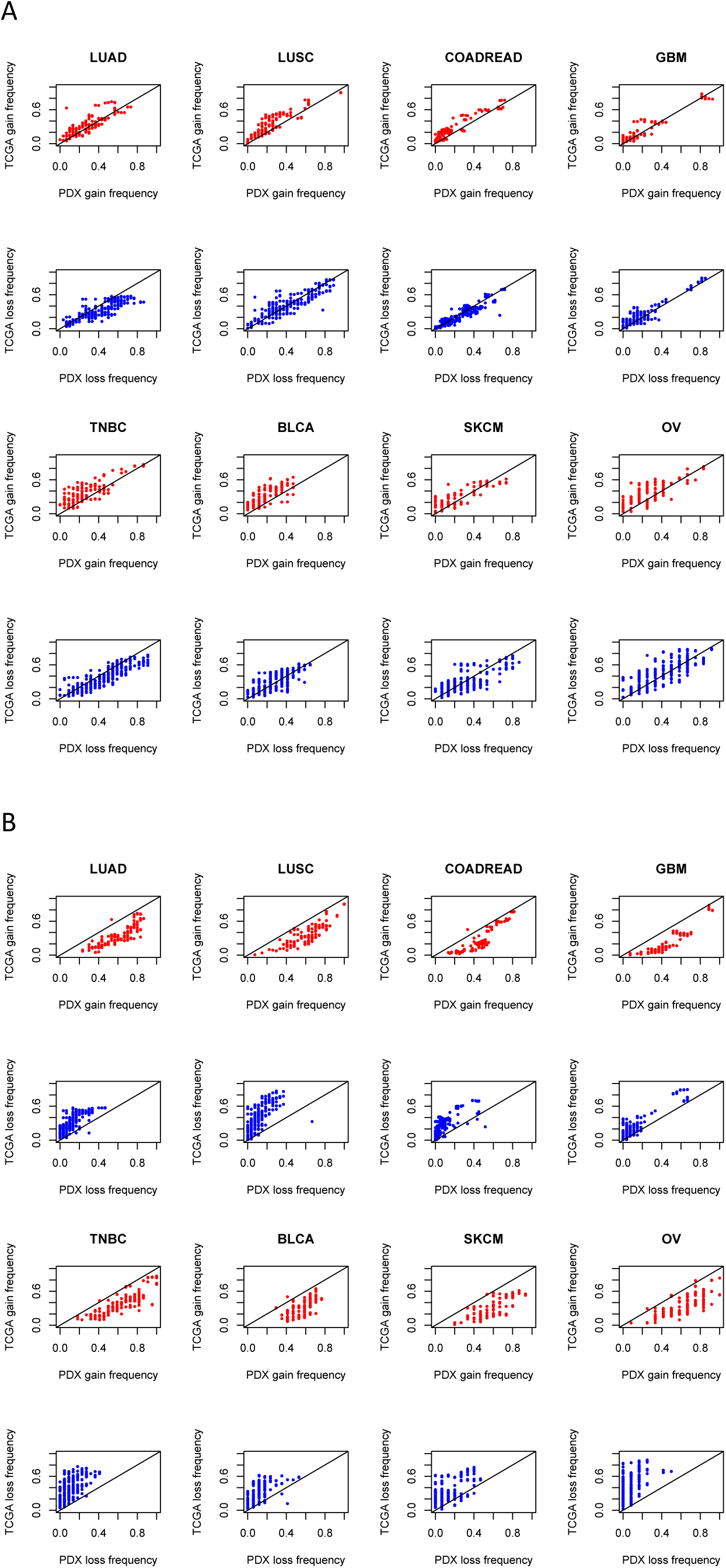
(A) Comparison of frequency of copy number gain (red) or loss (blue) of selected genes frequently amplified or deleted in TCGA tumors predicted by GISTIC analysis for each tumor type between PDX and TCGA datasets using predicted ploidy as a reference state. (B) Comparison of frequency of copy number gain (red) or loss (blue) of genes frequently amplified or deleted in TCGA tumors predicted by GISTIC analysis for each tumor type between PDX and TCGA datasets using diploid as a reference state.

**Figure S15.**
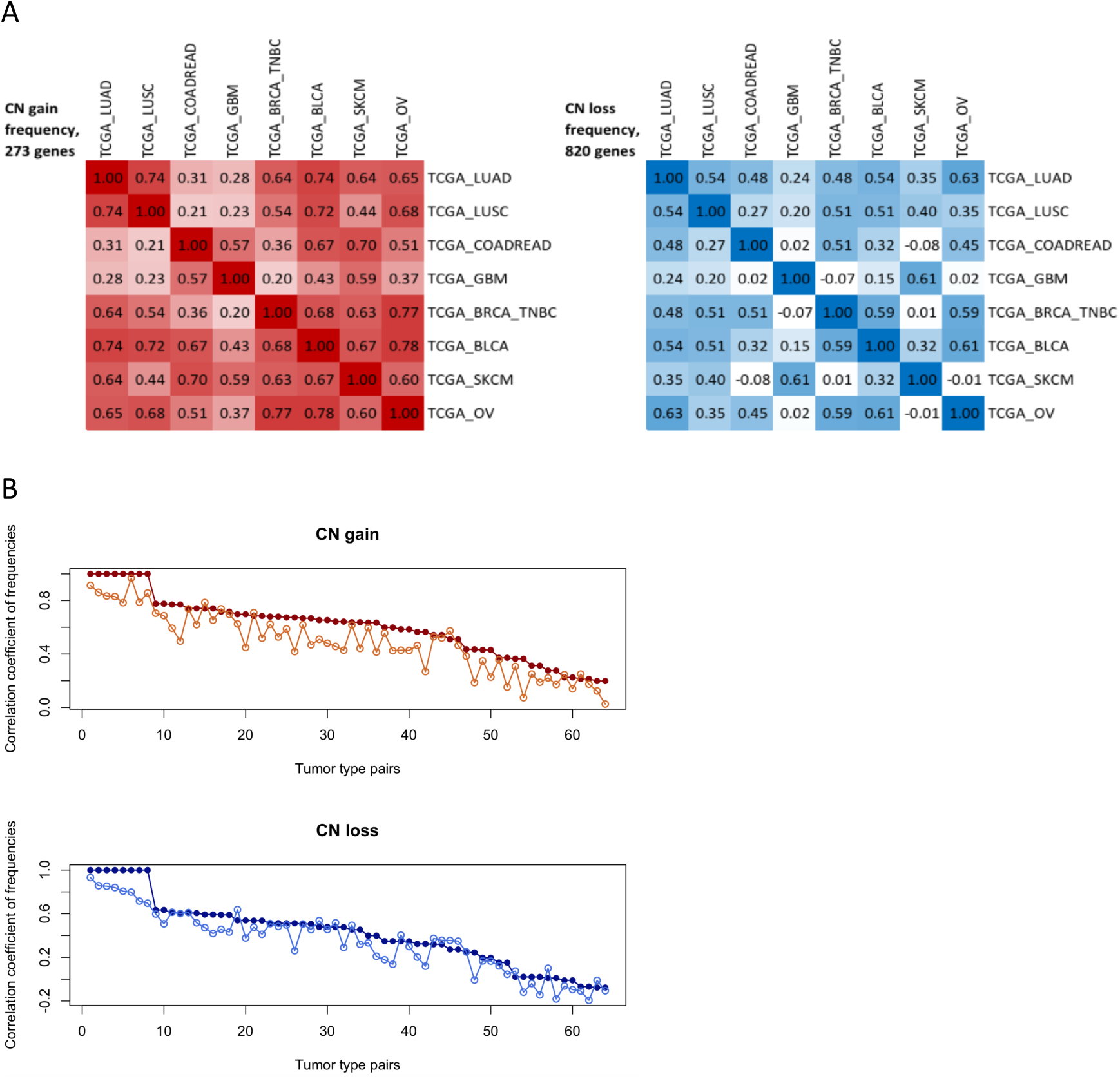
(A) Correlation of frequency of copy number gain (red) or loss (blue) of selected genes frequently amplified or deleted in TCGA tumors predicted by GISTIC analysis between each tumor type in TCGA SNP array datasets. (B) Ranked correlation coefficients based on Figure 4b and Supplementary Figure S6.

**Figure S16.**
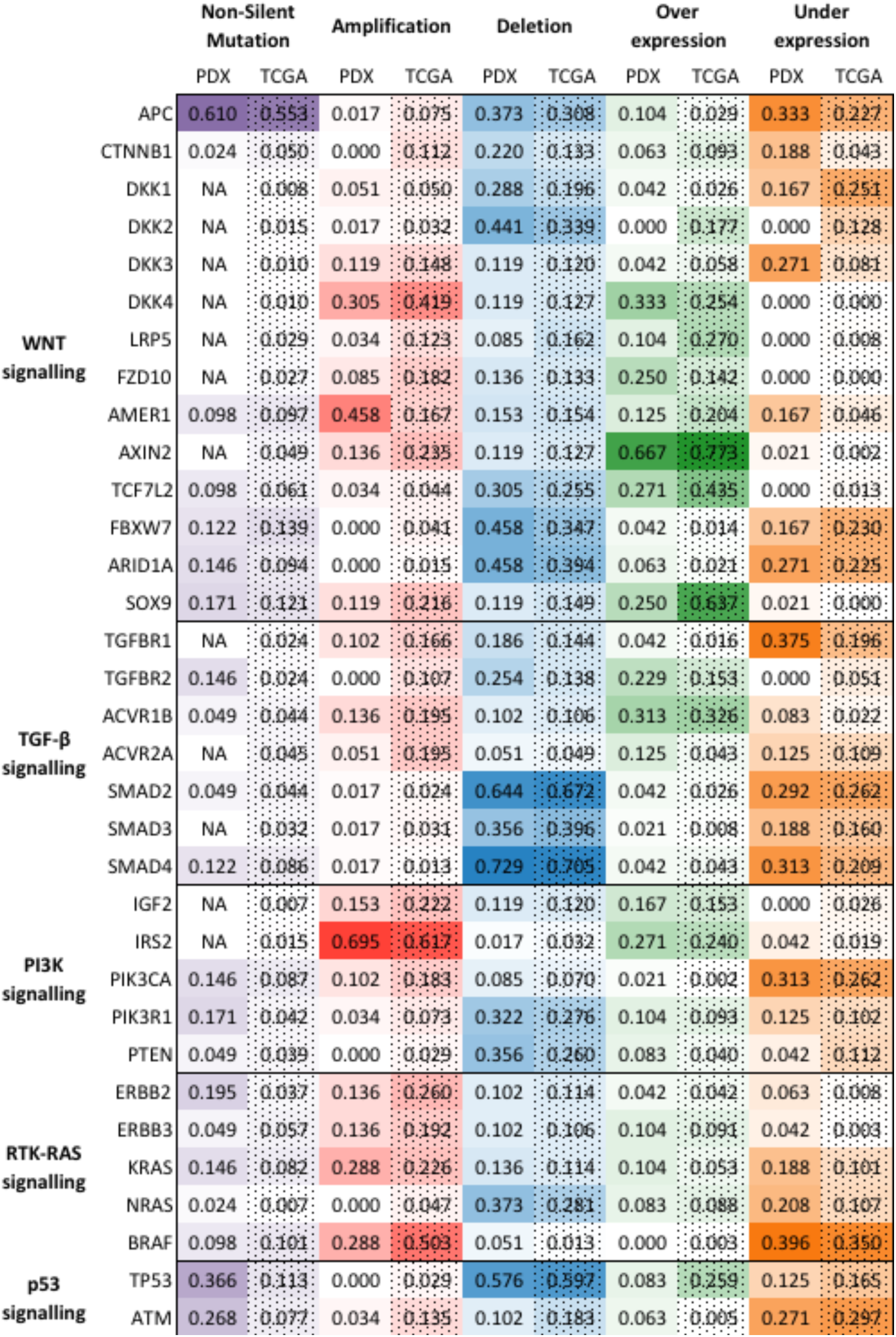
Frequency of genes altered PDX and TCGA tumors for each genomic datatype for colorectal cancer. These genes are identified by commonly affected pathways in colorectal cancer reported in TCGA studies. For both PDX and TCGA cohorts of colorectal cancer, we observed high frequencies in the 1) mutation of APC and over-expression of AXIN2 in the WNT signaling pathway, 2) amplification of IRS2 in the PI3K signaling pathway, 3) copy number loss of SMAD2 and SMAD4 in the TGF-β signaling pathway, 4) under-expression of BRAF in the RTK-RAS signaling pathway, and 5) copy number loss of TP53 in the TP53 signaling pathway.

**Supplementary Table S1.**
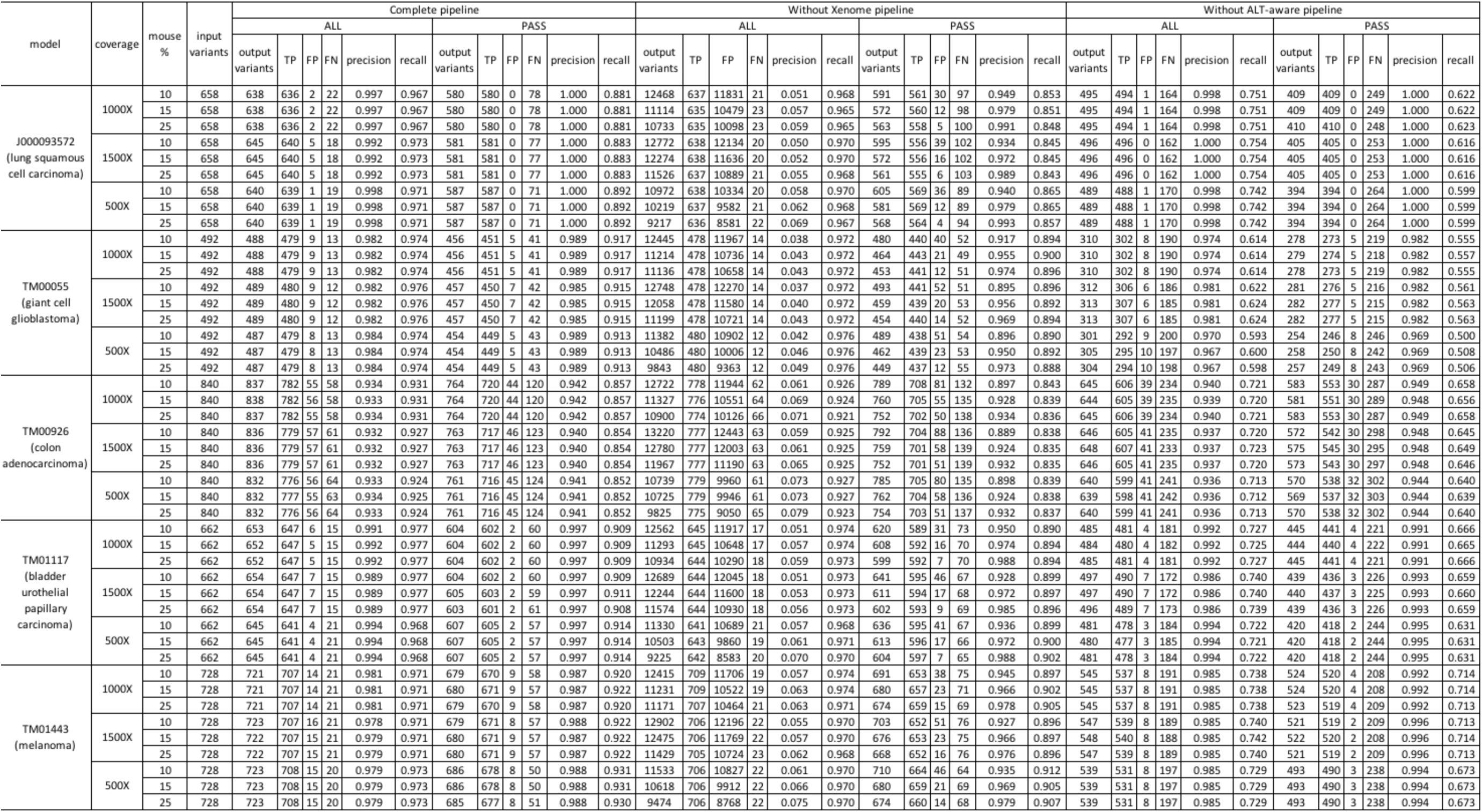
This table summarizes the results from the benchmarking studies of the CTP variant calling pipeline using 45 simulated sequencing datasets different samples, sequencing coverages, and mouse DNA content.

**Supplementary Table S2.** This table shows the correlation and difference in median of alternate allele frequencies between input and true positive variants for all the simulated samples. ALL: all variants called by the pipeline; PASS: variants annotated as “PASS” in the pipeline which pass the hard filters, minimum read depth and minimum alternate allele frequency of the variant.

| Model | Coverage (X) | Mouse percent | ALT_AF Correlation |  |  |  |  |  | ALT_AF Difference Median |  |  |  |  |  |
| --- | --- | --- | --- | --- | --- | --- | --- | --- | --- | --- | --- | --- | --- | --- |
|  |  |  | Output ALL | Output PASS | NoXenome ALL | NoXenome PASS | NoAltAware ALL | NoAltAware PASS | Output ALL | Output PASS | NoXenome ALL | NoXenome PASS | NoAltAware ALL | NoAltAware PASS |
| J000093572 | 500 | 10 | 0.995 | 0.995 | 0.985 | 0.984 | 0.982 | 0.981 | -3 | -3 | -3 | -3 | -13 | -15 |
|  |  | 15 | 0.995 | 0.995 | 0.976 | 0.975 | 0.982 | 0.981 | -3 | -3 | -3 | -3 | -13 | -15 |
|  |  | 25 | 0.995 | 0.995 | 0.957 | 0.952 | 0.982 | 0.981 | -3 | -3 | -3 | -3 | -13 | -15 |
|  | 1000 | 10 | 0.996 | 0.996 | 0.986 | 0.985 | 0.983 | 0.984 | -3 | -3 | -3 | -3 | -13.5 | -14 |
|  |  | 15 | 0.996 | 0.996 | 0.977 | 0.974 | 0.983 | 0.984 | -3 | -3 | -3 | -3 | -14 | -15 |
|  |  | 25 | 0.996 | 0.996 | 0.960 | 0.955 | 0.983 | 0.984 | -3 | -3 | -3 | -3 | -13 | -15 |
|  | 1500 | 10 | 0.997 | 0.997 | 0.986 | 0.985 | 0.976 | 0.974 | -3 | -3 | -3 | -3 | -13 | -15 |
|  |  | 15 | 0.997 | 0.997 | 0.977 | 0.974 | 0.976 | 0.974 | -3 | -3 | -3 | -3 | -13 | -15 |
|  |  | 25 | 0.997 | 0.997 | 0.961 | 0.955 | 0.976 | 0.974 | -3 | -3 | -3 | -3 | -13 | -15 |
| TM00055 | 500 | 10 | 0.991 | 0.990 | 0.954 | 0.927 | 0.970 | 0.958 | -3 | -3 | -4 | -4 | -12 | -12 |
|  |  | 15 | 0.990 | 0.990 | 0.931 | 0.878 | 0.970 | 0.958 | -3 | -3 | -4 | -4 | -12 | -12 |
|  |  | 25 | 0.990 | 0.990 | 0.888 | 0.777 | 0.970 | 0.959 | -3 | -3 | -4 | -4 | -12 | -12 |
|  | 1000 | 10 | 0.990 | 0.989 | 0.952 | 0.935 | 0.966 | 0.955 | -3 | -3 | -4 | -4 | -12 | -12 |
|  |  | 15 | 0.991 | 0.989 | 0.929 | 0.894 | 0.963 | 0.950 | -3 | -3 | -4 | -4 | -12 | -12 |
|  |  | 25 | 0.991 | 0.989 | 0.888 | 0.813 | 0.963 | 0.951 | -3 | -3 | -4 | -4 | -12 | -12 |
|  | 1500 | 10 | 0.991 | 0.988 | 0.953 | 0.931 | 0.968 | 0.953 | -3 | -4 | -4 | -4 | -12 | -12 |
|  |  | 15 | 0.991 | 0.989 | 0.930 | 0.882 | 0.969 | 0.954 | -3 | -4 | -4 | -4 | -12 | -12 |
|  |  | 25 | 0.991 | 0.989 | 0.888 | 0.811 | 0.969 | 0.953 | -3 | -4 | -4 | -4 | -12 | -12 |
| TM00926 | 500 | 10 | 0.990 | 0.990 | 0.974 | 0.977 | 0.976 | 0.980 | -3 | -3 | -3 | -3 | -14 | -15 |
|  |  | 15 | 0.990 | 0.990 | 0.963 | 0.965 | 0.976 | 0.980 | -3 | -3 | -3 | -3 | -14 | -15 |
|  |  | 25 | 0.990 | 0.990 | 0.939 | 0.943 | 0.976 | 0.980 | -3 | -3 | -3 | -3 | -14 | -15 |
|  | 1000 | 10 | 0.992 | 0.992 | 0.977 | 0.980 | 0.977 | 0.982 | -3 | -3 | -3 | -3 | -15 | -15 |
|  |  | 15 | 0.992 | 0.992 | 0.965 | 0.970 | 0.977 | 0.982 | -3 | -3 | -3 | -3 | -15 | -15 |
|  |  | 25 | 0.992 | 0.992 | 0.943 | 0.947 | 0.977 | 0.982 | -3 | -3 | -3 | -3 | -15 | -15 |
|  | 1500 | 10 | 0.993 | 0.993 | 0.977 | 0.981 | 0.974 | 0.976 | -3 | -3 | -3 | -3 | -14 | -15 |
|  |  | 15 | 0.992 | 0.993 | 0.966 | 0.971 | 0.974 | 0.976 | -3 | -3 | -3 | -3 | -14 | -15 |
|  |  | 25 | 0.992 | 0.993 | 0.945 | 0.950 | 0.974 | 0.977 | -3 | -3 | -3 | -3 | -14 | -15 |
| TM01117 | 500 | 10 | 0.994 | 0.992 | 0.979 | 0.979 | 0.977 | 0.977 | -3 | -3 | -3 | -3 | -13 | -14 |
|  |  | 15 | 0.994 | 0.993 | 0.969 | 0.968 | 0.977 | 0.977 | -3 | -3 | -3 | -3 | -13 | -14 |
|  |  | 25 | 0.994 | 0.992 | 0.946 | 0.942 | 0.977 | 0.977 | -3 | -3 | -3 | -3 | -13 | -14 |
|  | 1000 | 10 | 0.995 | 0.994 | 0.979 | 0.981 | 0.981 | 0.983 | -3 | -3 | -3 | -3 | -14 | -14 |
|  |  | 15 | 0.995 | 0.994 | 0.969 | 0.969 | 0.981 | 0.983 | -3 | -3 | -3 | -3 | -14 | -14 |
|  |  | 25 | 0.995 | 0.994 | 0.949 | 0.945 | 0.981 | 0.983 | -3 | -3 | -3 | -3 | -14 | -14 |
|  | 1500 | 10 | 0.995 | 0.995 | 0.980 | 0.982 | 0.978 | 0.978 | -3 | -3 | -3 | -3 | -13 | -14 |
|  |  | 15 | 0.995 | 0.994 | 0.971 | 0.971 | 0.979 | 0.979 | -3 | -3 | -3 | -3 | -13 | -13.5 |
|  |  | 25 | 0.995 | 0.995 | 0.951 | 0.947 | 0.978 | 0.977 | -3 | -3 | -3 | -3 | -13 | -14 |
| TM01443 | 500 | 10 | 0.984 | 0.988 | 0.962 | 0.970 | 0.971 | 0.972 | -3 | -3 | -3 | -3 | -14 | -15 |
|  |  | 15 | 0.984 | 0.988 | 0.950 | 0.959 | 0.971 | 0.972 | -3 | -3 | -3 | -3 | -14 | -15 |
|  |  | 25 | 0.984 | 0.988 | 0.921 | 0.934 | 0.971 | 0.972 | -3 | -3 | -4 | -4 | -14 | -15 |
|  | 1000 | 10 | 0.981 | 0.990 | 0.961 | 0.973 | 0.970 | 0.972 | -3 | -3 | -4 | -3 | -15 | -15 |
|  |  | 15 | 0.982 | 0.990 | 0.947 | 0.959 | 0.970 | 0.972 | -3 | -3 | -4 | -4 | -15 | -15 |
|  |  | 25 | 0.982 | 0.990 | 0.925 | 0.937 | 0.970 | 0.972 | -3 | -3 | -4 | -4 | -15 | -15 |
|  | 1500 | 10 | 0.984 | 0.991 | 0.962 | 0.974 | 0.968 | 0.968 | -3 | -3 | -3 | -3 | -14 | -15 |
|  |  | 15 | 0.984 | 0.991 | 0.949 | 0.960 | 0.967 | 0.967 | -3 | -3 | -3 | -4 | -15 | -15 |
|  |  | 25 | 0.984 | 0.991 | 0.924 | 0.939 | 0.968 | 0.967 | -3 | -3 | -4 | -4 | -14 | -15 |

**Supplementary Table S3.**
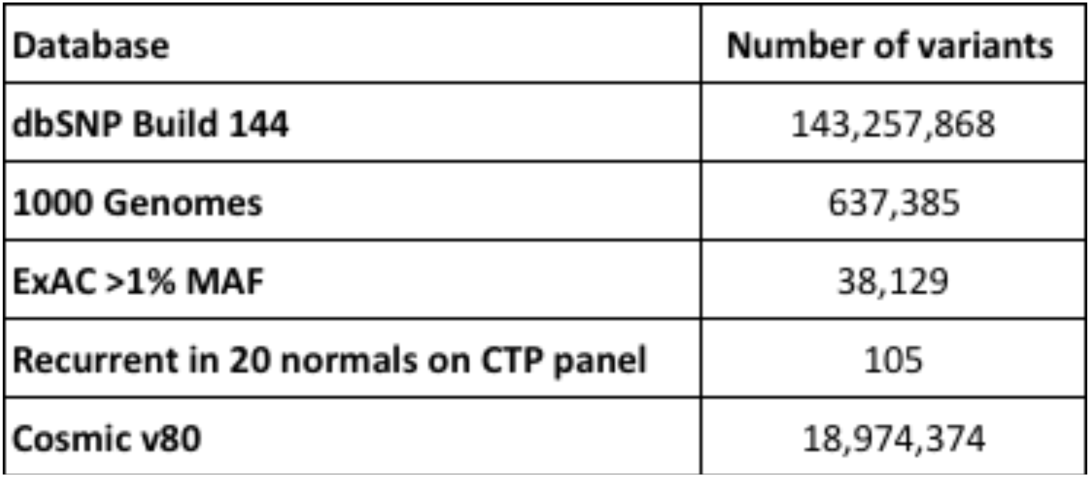
Number of variants in each germline databases.

| Database | Number of variants |
| --- | --- |
| dbSNP Build 144 | 143,257,868 |
| 1000 Genomes | 637,385 |
| ExAC >1% MAF | 38,129 |
| Recurrent in 20 normals on CTP panel | 105 |
| Cosmic v80 | 18,974,374 |

**Supplementary Table S4.** Number of unique variants in each CTP sample (n=383), represented by mean, median and standard deviation, called by GATK and after each filtering or rescue step. The last row shows the average number of variants annotated as clinically relevant.

| CTP (383 samples) | SNP |  |  | INDEL |  |  |
| --- | --- | --- | --- | --- | --- | --- |
| Number of unique variants | Mean | Median | Standard deviation | Mean | Median | Standard deviation |
| Total | 798 | 779 | 107 | 30 | 28 | 13 |
| DP, AF and Hard filters | 601 | 592 | 80 | 23 | 22 | 11 |
| Germline filters | 164 | 146 | 84 | 9.9 | 9 | 7.9 |
| Recurrent variants filter and High/Moderate impact variants | 67 | 57 | 42 | 3.1 | 2 | 3.2 |
| Clinically relevant variants rescue | 71 | 61 | 42 | 3.2 | 2 | 3.6 |
| Clinically relevant variants | 5.6 | 5 | 2.1 | 0.18 | 0 | 0.65 |

**Supplementary Table S5.**
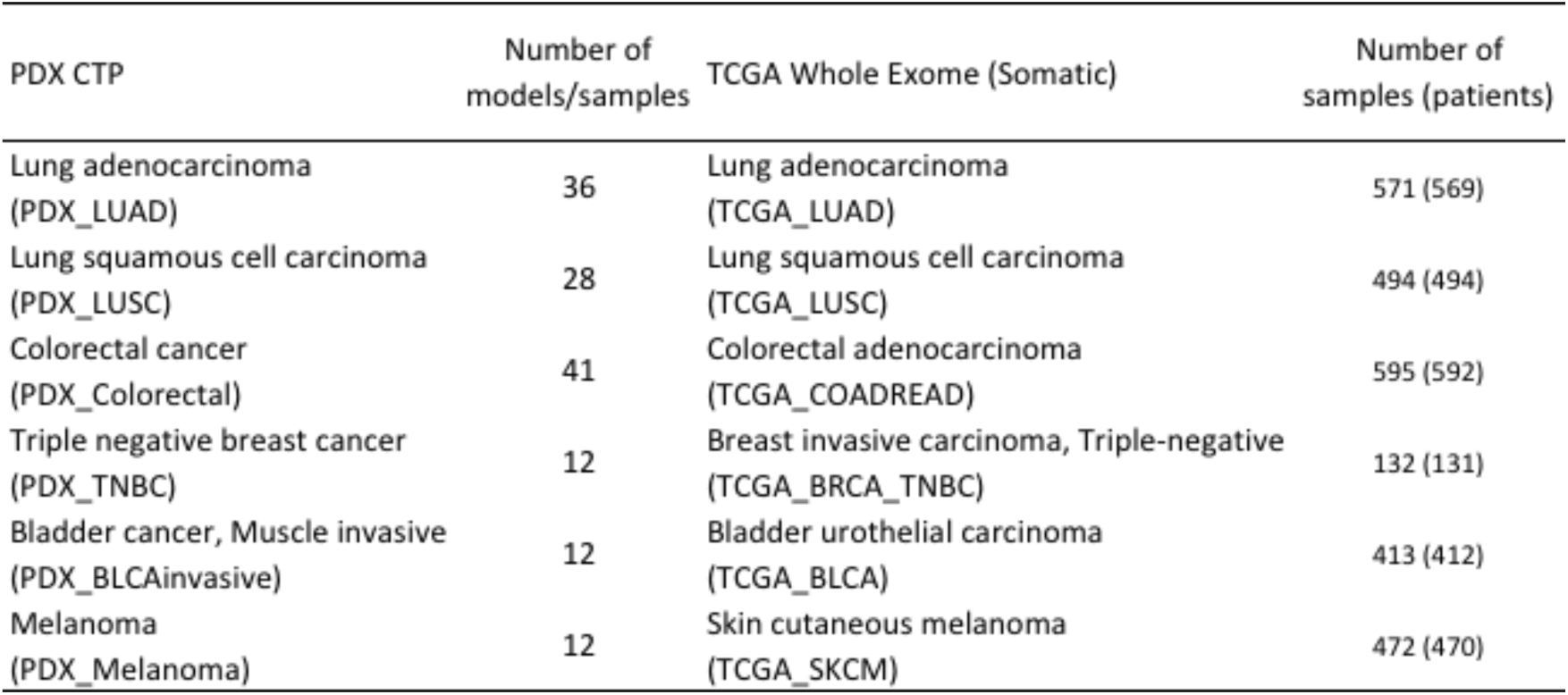
Number of PDX and TCGA samples for 5 tumor types used for analysis of variant calling.

**Supplementary Table S6.** Classifier gene table to classify EBV-associated PDX lymphomas versus other tumors. Up-regulation: +1; Down-regulation: −1.

| Ensembl ID | Gene Symbol | Up or Down regulation |
| --- | --- | --- |
| ENSG00000002919 | SNX11 | 1 |
| ENSG00000007866 | TEAD3 | -1 |
| ENSG00000009790 | TRAF3IP3 | 1 |
| ENSG00000028277 | POU2F2 | 1 |
| ENSG00000056558 | TRAF1 | 1 |
| ENSG00000061676 | NCKAP1 | -1 |
| ENSG00000062370 | ZFP112 | -1 |
| ENSG00000072818 | ACAP1 | 1 |
| ENSG00000076662 | ICAM3 | 1 |
| ENSG00000084070 | SMAP2 | 1 |
| ENSG00000086730 | LAT2 | 1 |
| ENSG00000088256 | GNA11 | -1 |
| ENSG00000102096 | PIM2 | 1 |
| ENSG00000104067 | TJP1 | -1 |
| ENSG00000104814 | MAP4K1 | 1 |
| ENSG00000110031 | LPXN | 1 |
| ENSG00000110777 | POU2AF1 | 1 |
| ENSG00000116473 | RAP1A | 1 |
| ENSG00000120256 | LRP11 | -1 |
| ENSG00000122386 | ZNF205 | -1 |
| ENSG00000126822 | PLEKHG3 | -1 |
| ENSG00000130147 | SH3BP4 | -1 |
| ENSG00000137693 | YAP1 | -1 |
| ENSG00000138185 | ENTPD1 | 1 |
| ENSG00000142192 | APP | -1 |
| ENSG00000144677 | CTDSPL | -1 |
| ENSG00000147799 | ARHGAP39 | -1 |
| ENSG00000150760 | DOCK1 | -1 |
| ENSG00000152990 | GPR125 | -1 |
| ENSG00000156052 | GNAQ | -1 |
| ENSG00000157985 | AGAP1 | -1 |
| ENSG00000162627 | SNX7 | -1 |
| ENSG00000163625 | WDFY3 | -1 |
| ENSG00000167984 | NLRC3 | 1 |
| ENSG00000172578 | KLHL6 | 1 |
| ENSG00000173200 | PARP15 | 1 |
| ENSG00000180096 | 1-Sep | 1 |
| ENSG00000180891 | CUEDC1 | -1 |
| ENSG00000187079 | TEAD1 | -1 |
| ENSG00000187164 | KIAA1598 | -1 |
| ENSG00000188822 | CNR2 | 1 |
| ENSG00000197702 | PARVA | -1 |
| ENSG00000197763 | TXNRD3 | -1 |
| ENSG00000198833 | UBE2J1 | 1 |
| ENSG00000211895 | IGHA1 | 1 |
| ENSG00000213402 | PTPRCAP | 1 |
| ENSG00000213999 | MEF2B | 1 |
| ENSG00000249096 | RP11-290F5.1 | 1 |

**Supplementary Table S7.** Pearson correlation coefficient of gene-based log2(total CN/ploidy) between the single-tumor and tumor-normal CNV analysis.

| Model ID | Sample ID | Correlation coefficient (Pearson) |
| --- | --- | --- |
| J000077712 | J000077712PT | 0.811 |
| J000079689 | J000095142 | 0.938 |
| J000079689 | J000079689PT | 0.983 |
| TM00327 | OVJX01F000P0 | 0.974 |
| TM00327 | OVJX01F017P0 | 0.955 |
| TM00327 | OVJX01F020P0 | 0.969 |
| TM00327 | OVJX01F021P0 | 0.968 |
| TM01594 | TM01594F062P2 | 0.966 |
| TM01594 | TM01594FPT | 0.978 |

**Supplementary Table S8.**
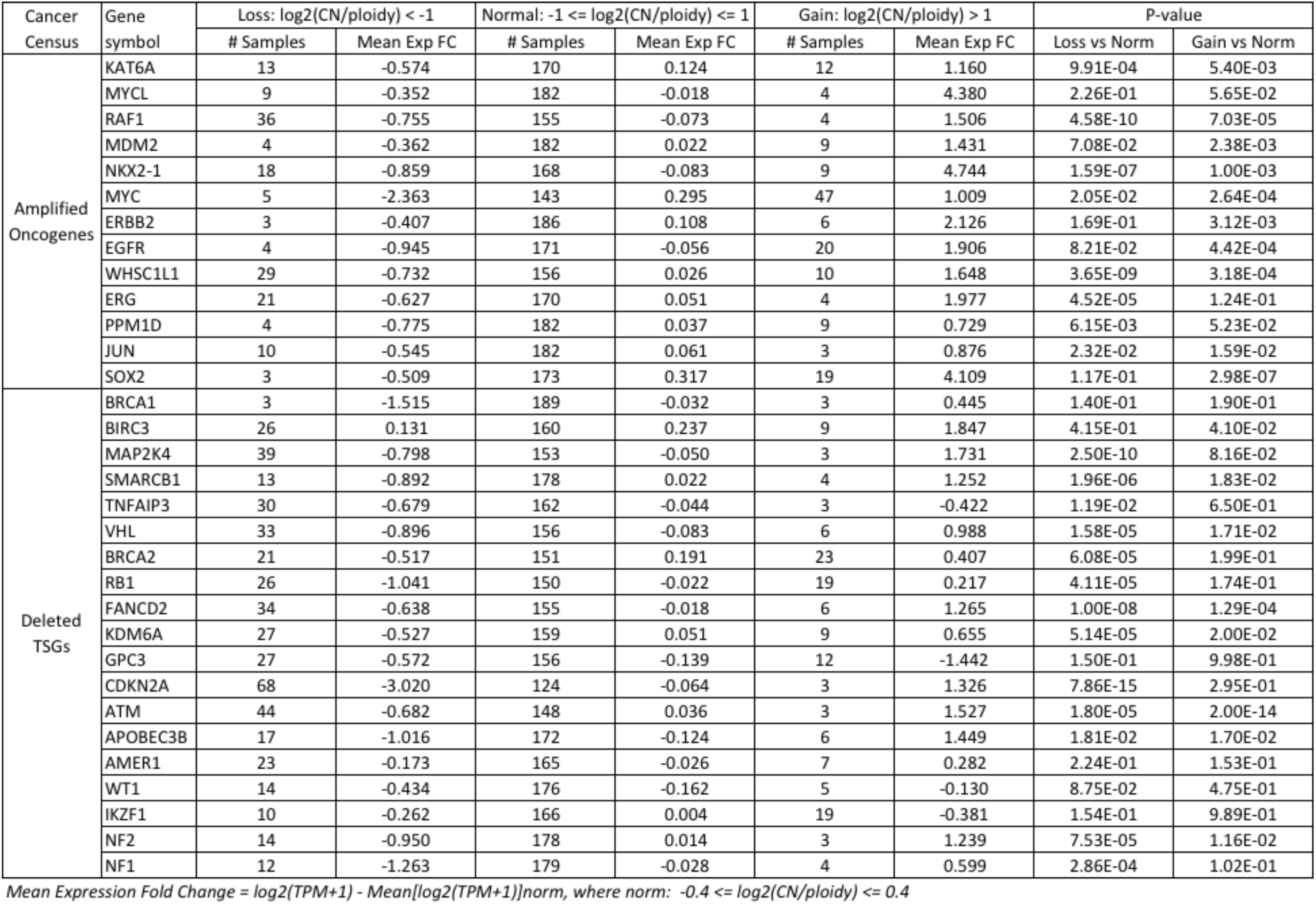
Mean expression fold change of genes with copy number normal, gain and loss state for genes found in the Cancer Census that is listed as oncogenes affected by amplification and tumor suppressor genes affected by deletions (refer to Supplementary Figure S16). The p-value, calculated by Student’s t-test, measures if the difference in expression for each gene between the copy number loss models versus normal models, and between the copy number gain models versus normal models is significant.

**Supplementary Table S9.** Contingency table for Fisher’s Exact Test for CTP genes with coding, non-silent mutations for different tumor types in the PDX and TCGA cohort. For PDX, the number of CTP genes with and without coding, non-silent mutations in each tumor type cohort was counted. For TCGA data with more samples than PDX, the number of CTP genes with coding, non-silent mutations at ≥5% and <5% frequency in each tumor type cohort was counted.

| COADREAD CTP (coding, non-silent) | TCGA mut $\geq 5\%$ | TCGA mut $< 5\%$ | Total |
| --- | --- | --- | --- |
| PDX mut | 54 | 244 | 298 |
| PDX not mut | 0 | 60 | 60 |
| Total | 54 | 304 | 358 |

| LUAD CTP (coding, non-silent) | TCGA mut $\geq 5\%$ | TCGA mut $< 5\%$ | Total |
| --- | --- | --- | --- |
| PDX mut | 44 | 220 | 264 |
| PDX not mut | 1 | 93 | 94 |
| Total | 45 | 313 | 358 |

| LUSC CTP (coding, non-silent) | TCGA mut $\geq 5\%$ | TCGA mut $< 5\%$ | Total |
| --- | --- | --- | --- |
| PDX mut | 29 | 197 | 226 |
| PDX not mut | 1 | 131 | 132 |
| Total | 30 | 328 | 358 |

| BLCA CTP (coding, non-silent) | TCGA mut $\geq 5\%$ | TCGA mut $< 5\%$ | Total |
| --- | --- | --- | --- |
| PDX mut | 37 | 151 | 188 |
| PDX not mut | 6 | 164 | 170 |
| Total | 43 | 315 | 358 |

| SKCM CTP (coding, non-silent) | TCGA mut $\geq 5\%$ | TCGA mut $< 5\%$ | Total |
| --- | --- | --- | --- |
| PDX mut | 84 | 145 | 229 |
| PDX not mut | 5 | 124 | 129 |
| Total | 89 | 269 | 358 |

| BRCA TNBC CTP (coding, non-silent) | TCGA mut $\geq 5\%$ | TCGA mut $< 5\%$ | Total |
| --- | --- | --- | --- |
| PDX mut | 18 | 173 | 191 |
| PDX not mut | 1 | 166 | 167 |
| Total | 19 | 339 | 358 |

**Supplementary Table S10.**
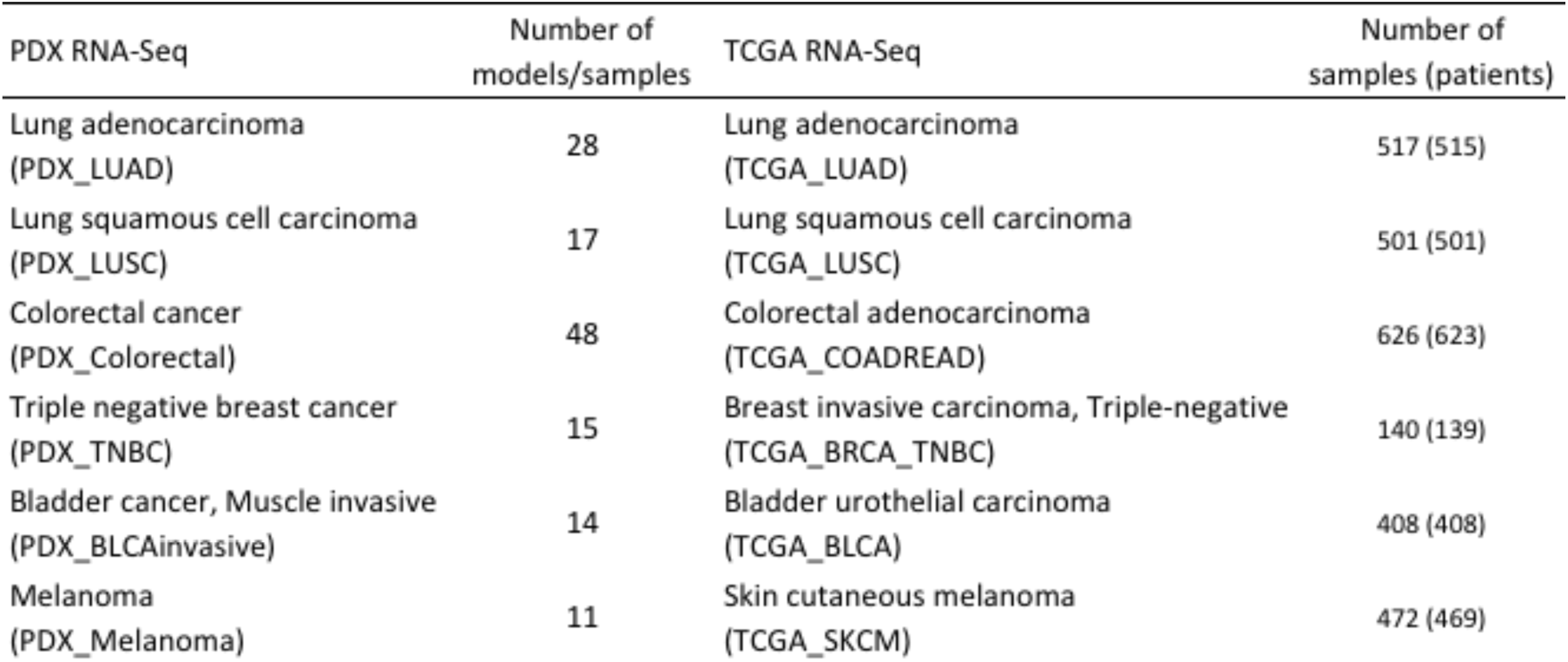
Number of PDX and TCGA samples for 6 tumor types used for analysis of RNA-Seq expression profiling.

**Supplementary Table S11.**
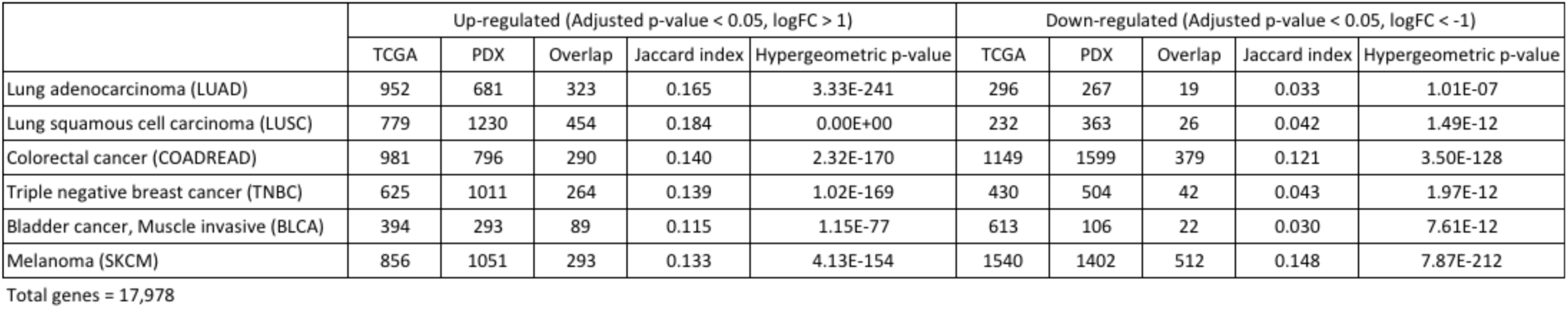
Number of genes that are up-regulated (adjusted p-value < 0.05, log (fold change of TPM+1) > 1 by limma) or down-regulated (adjusted p-value < 0.05, log (fold change of TPM+1) < −1 by limma) for each tumor types versus all other tumor types for PDX and TCGA RNA-Seq data respectively. This table shows the overlap of each set of genes between PDX and TCGA RNA-Seq data.

**Supplementary Table S12.** Number of PDX and TCGA samples for 8 tumor types used for analysis of copy number and LOH predicted from SNP array.

| PDX CNV | Number of models/samples | TCGA CNV | Number of tumors/samples |
| --- | --- | --- | --- |
| Lung adenocarcinoma (PDX_LUAD) | 30 | Lung adenocarcinoma (TCGA_LUAD) | 516 |
| Lung squamous cell carcinoma (PDX_LUSC) | 27 | Lung squamous cell carcinoma (TCGA_LUSC) | 501 |
| Colorectal cancer (PDX_Colorectal) | 60 | Colorectal adenocarcinoma (TCGA_COADREAD) | 616 |
| Glioblastoma multiforme (PDX_GBM) | 27 | Glioblastoma multiforme (TCGA_GBM) | 577 |
| Triple negative breast cancer (PDX_TNBC) | 22 | Breast invasive carcinoma, Triple-negative (TCGA_BRCA_TNBC) | 136 |
| Bladder cancer, Muscle invasive (PDX_BLCAinvasive) | 17 | Bladder urothelial carcinoma (TCGA_BLCA) | 408 |
| Melanoma (PDX_Melanoma) | 15 | Skin cutaneous melanoma (TCGA_SKCM) | 367 |
| Ovarian carcinoma (PDX_OVcarcinoma) | 12 | Ovarian serous cystadenocarcinoma (TCGA_OV) | 579 |

